# Multivalent proteins rapidly and reversibly phase-separate upon osmotic cell volume change

**DOI:** 10.1101/748293

**Authors:** Ameya P. Jalihal, Sethuramasundaram Pitchiaya, Lanbo Xiao, Pushpinder Bawa, Xia Jiang, Karan Bedi, Abhijit Parolia, Marcin Cieslik, Mats Ljungman, Arul M. Chinnaiyan, Nils G. Walter

## Abstract

Processing bodies (PBs) and stress granules (SGs) are prominent examples of sub-cellular, membrane-less compartments that are observed under physiological and stress conditions, respectively. We observe that the trimeric PB protein DCP1A rapidly (within ∼10 s) phase-separates in mammalian cells during hyperosmotic stress and dissolves upon isosmotic rescue (over ∼100 s) with minimal impact on cell viability even after multiple cycles of osmotic perturbation. Strikingly, this rapid intracellular hyperosmotic phase separation (HOPS) correlates with the degree of cell volume compression, distinct from SG assembly, and is exhibited broadly by homo-multimeric (valency ≥ 2) proteins across several cell types. Notably, HOPS sequesters pre-mRNA cleavage factor components from actively transcribing genomic loci, providing a mechanism for hyperosmolarity-induced global impairment of transcription termination. Together, our data suggest that the multimeric proteome rapidly responds to changes in hydration and molecular crowding, revealing an unexpected mode of globally programmed phase separation and sequestration that adapts the cell to volume change.

**GRAPHICAL ABSTRACT:** 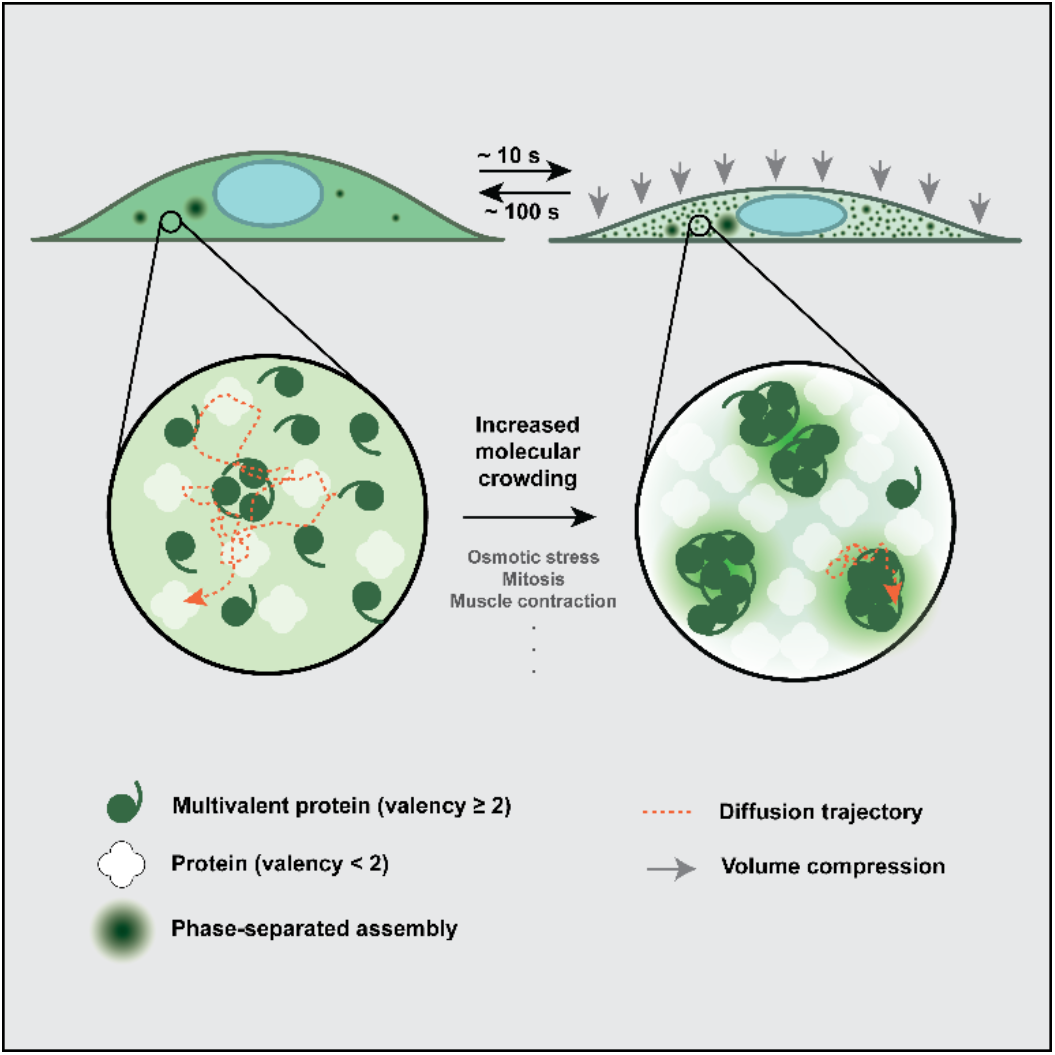

**IN BRIEF:** Cells constantly experience osmotic variation. These external changes lead to changes in cell volume, and consequently the internal state of molecular crowding. Here, Jalihal and Pitchiaya et al. show that multimeric proteins respond rapidly to such cellular changes by undergoing rapid and reversible phase separation.

**HIGHLIGHTS:**

- DCP1A undergoes rapid and reversible hyperosmotic phase separation (HOPS)
- HOPS of DCP1A depends on its trimerization domain
- Self-interacting multivalent proteins (valency ≥ 2) undergo HOPS
- HOPS of CPSF6 explains transcription termination defects during osmotic stress

## INTRODUCTION

Membrane-less condensates, often referred to as membrane-less organelles (MLOs), represent sub-cellular sites within the cytosol or nucleus of mammalian cells, wherein key processes such as transcription, translation, post-transcriptional gene regulation, and metabolism are altered compared to the nucleoplasmic or cytoplasmic bulk (Banani et al., 2017; Spector, 2006). Mis-regulation of MLOs and the *de novo* condensation of mutated proteins into MLOs have been strongly associated with altered gene regulation (Berchtold et al., 2018) and severe pathologies such as amyotrophic lateral sclerosis (ALS) (Banani et al., 2017; Patel et al., 2015; Shin and Brangwynne, 2017). Therefore, understanding the cellular mechanisms by which these structures assemble should yield insights critical for understanding both cellular physiology and disease (Alberti, 2017; Hyman et al., 2014; Toretsky and Wright, 2014).

MLOs are hypothesized to arise from the phase separation of dispersed multivalent biomolecules under specific conditions of pH, temperature, and concentration (Boeynaems et al., 2018; Hyman et al., 2014; Shin and Brangwynne, 2017). Extensive evaluation of this notion *in vitro* has defined the molecular features required to form MLOs (Hyman et al., 2014; Shin and Brangwynne, 2017; Wang et al., 2018), especially in the context of homotypic or heterotypic interactions of low complexity domain (LCD) containing proteins and RNAs, and has yielded an ever-expanding list of cellular components that can spontaneously phase-separate in the test tube. Yet, the significance of the propensity of these biomolecules to phase-separate under the physiological conditions of their native intracellular environment, where molecular crowding is dominant, is poorly understood (Alberti et al., 2019). While it is possible to alter crowding within the test tube via the addition of synthetic macromolecules (Alberti et al., 2019), the nature and extent of crowding in the cellular context is quite different (Daher et al., 2018; Walter, 2019) and dynamically changes with the cellular state. For example, cell volume adjustments occur during processes critical to both cellular homeostasis and pathology, including the cell cycle (Tzur et al., 2009; Zlotek-Zlotkiewicz et al., 2015) as well as upon cell adhesion and migration (Guo et al., 2017; Watkins and Sontheimer, 2011). Changes in cell volume and molecular crowding, frequently encountered by cells of the kidney, liver, and gut (Lang et al., 1998), are even more rapid and dramatic during osmotic perturbation (Guo et al., 2017; Hersen et al., 2008; Miermont et al., 2013). How cells respond to rapid and frequent volume perturbations with seemingly minimal impact on their viability and whether the resulting dynamic changes in macromolecular crowding affect intracellular phase separation remain unknown.

Processing bodies (PBs) are an example of gene regulatory MLOs that are constitutively present in eukaryotic cells under physiological conditions and endogenous concentrations of the constituents (Anderson and Kedersha, 2009). Their intracellular copy number has been shown to be modulated not only during the cell cycle (Aizer et al., 2013), but also upon prolonged (minutes to hours) hypertonic or hyperosmotic stress (Huch and Nissan, 2017), conditions which can lead to nephritic and vascular pathologies (Brocker et al., 2012). Much like other environmental stressors (e.g., heat shock, oxidative stress, metabolite deprivation), prolonged hyperosmotic stress also triggers the integrated stress response (ISR) and the formation of a type of gene-regulatory MLOs called stress granules (SGs) (Anderson and Kedersha, 2009). While both PBs and SGs are thought to assemble via a conceptually similar mechanism involving multivalent interactions between non-translating mRNAs and LCD-bearing RNA binding proteins (Van Treeck and Parker, 2018), they are compositionally distinct (Hubstenberger et al., 2017; Jain et al., 2016; Khong et al., 2017). Whether components of PBs and SGs are affected differentially by distinct stresses is largely unknown. Given that hyperosmotic stress rapidly (seconds to minutes) imparts cell volume change (Guo et al., 2017; Hersen et al., 2008; Miermont et al., 2013), it is also unclear whether the ISR, and consequently SGs, can be induced at this time scale. Finally, the observation that PBs are similarly regulated by the cell cycle and hypertonic stress (Aizer et al., 2013; Huch and Nissan, 2017) raises the question of whether PB regulation and cell volume change may be connected.

Here we investigate the role of macromolecular crowding and cell volume change on the intracellular phase separation of proteins using osmotic perturbations. We observe that DCP1A, a marker of PBs and component of the mRNA decapping machinery, rapidly (within ∼10 s) undergoes cytosolic phase separation in response to hypertonic stress and that these condensates dissolve over ∼100 s upon return to isotonic medium. This hyperosmotic phase separation (HOPS) can be cycled multiple times with minimal impact on cell viability, and is caused by changes in cellular water content and molecular crowding since its extent is directly proportional to the osmotic compression of the cell. We further find that HOPS is induced by DCP1A’s homo-trimerization domain and observed across a variety of cell types. Strikingly, most multimeric proteins tested with a valency of at least 2 (i.e., forming trimers and higher order multimers, but not dimers and monomers) undergo HOPS, strongly suggesting that rapid changes in hydration and molecular crowding are sensed by a significant fraction of the proteome and may lead to pleiotropic effects. Notably, G3BP and polyA RNA, as markers of SGs, do not undergo HOPS (as characterized by condensation within ∼10 s) at their endogenous concentrations, supporting the notion that it is a unique feature of multimeric proteins. HOPS of multimeric Cleavage and Polyadenylation Specific Factor 6 (CPSF6) within the cell’s nucleus is correlated with widespread impairment of transcription termination, apparently due to sequestration of the pre-mRNA cleavage complex from a subset of transcription end sites (TES). Our findings suggest that HOPS is a heretofore-underappreciated driver of protein phase separation that rapidly senses changes in cell volume with profound impact on cellular homeostasis.

## RESULTS

### Changes in extracellular tonicity induce rapid and reversible intracellular phase separation of DCP1A, but not SG markers

In a previous study (Pitchiaya et al., 2019), we observed that osmotic stress leads to phase separation of DCP1A, a non-catalytic protein component of the eukaryotic decapping complex and conserved PB marker (Anderson and Kedersha, 2009). To study the intracellular kinetics of de novo PB and SG formation in response to stress more broadly, we subjected U2OS cells to osmotic and oxidative stressors, and performed fixed-cell protein immunofluorescence (IF) or combined IF and RNA fluorescent *in situ* hybridization (RNA-FISH) before and after the stressors (Figure 1). Under isotonic conditions (150 mM Na^+^), ∼9% of cellular DCP1A localized within ∼10-30 foci (each ranging ∼300-800 nm in diameter) per cell, whereas G3BP protein and polyA RNA, as markers of SGs (Patel et al., 2015), were dispersed throughout the cytosol (Figures 1A and S1A). Upon a short (2 min) hypertonic (300 mM Na^+^) shock, ∼50% of cellular DCP1A, but neither G3BP nor polyA RNA, localized within ∼200-300 smaller (200-300 nm) foci per cell (Figures 1A and S1A). Moreover, IF-based colocalization analysis showed that these newly formed DCP1A foci do not contain other PB markers (e.g., EDC4 and DDX6, Figure S1B), suggesting that hypertonicity induced DCP1A foci are not bona fide PBs. No further change in DCP1A foci number or partitioning extent was observed even after prolonged (60 min) hypertonic treatment. At this later time point however, both G3BP and polyA RNA showed significant focus formation (∼100-200 foci per cell, 200-300 nm in diameter; Figures 1A and S1A), in line with previous observations (Bounedjah et al., 2012), with ∼13% of cellular G3BP partitioning into foci. By contrast, after 2 min of oxidative stress with sodium arsenite (SA), the number and partition coefficient of DCP1A foci were similar to unstressed cells, and G3BP and polyA RNA were still dispersed throughout the cytosol (Figures 1B and S1C). In this short time frame (2 min), phosphorylated EIF4E (P-EIF4E) mediated ISR was not induced by any of the stresses tested (Figure S1D). Upon prolonged (60 min) SA treatment, the number of DCP1A foci only marginally increased (∼25-40 foci per cell, 300-800 nm in diameter), with a concomitant small increase in partition coefficient (∼14%, Figures 1B and S1C). As expected for the induction of the ISR at these conditions, G3BP and polyA RNA formed a small number of (∼ 18% of cellular G3BP localized in ∼10-30 foci per cell) large (400-1100 nm in diameter) foci and P-EIF4E was induced (Figures 1B, S1C and S1D). Together, these data suggest that DCP1A and G3BP, when visualized in their physiological contexts, assemble into microscopically detectable foci at distinct rates and extents in response to osmotic and oxidative stressors.

**Figure 1.**
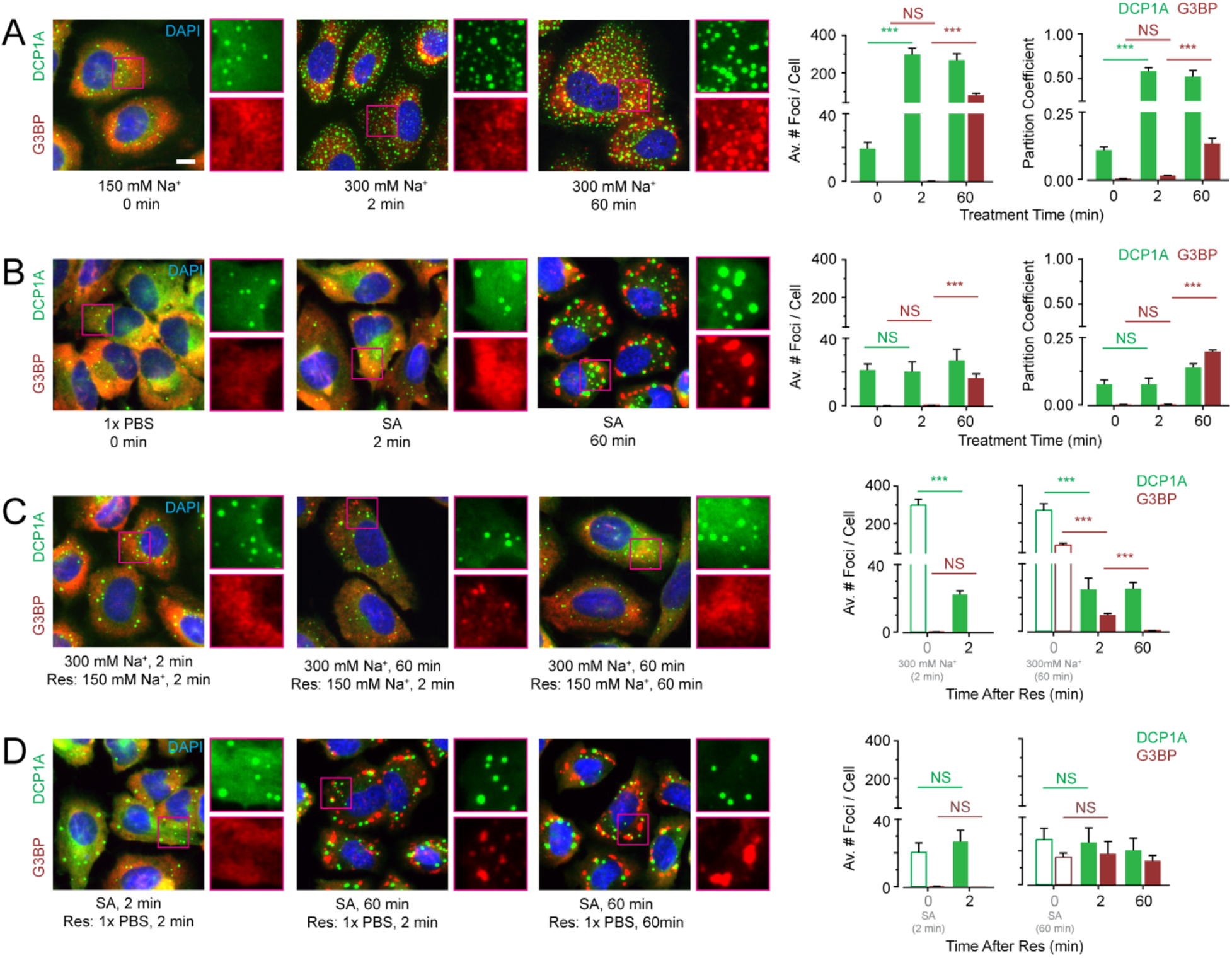
Extent and kinetics of DCP1A phase separation during hypertonic stress are distinct from those of SG markers G3BP. (A-D) Representative pseudocolored immunofluorescence (IF) images of U2OS cells stained for DAPI (blue), DCP1A (green) or G3BP (red) and the corresponding quantification of average number of spots per cell. Scale bar, 10 µm. (A) Cells were treated with isotonic (150 mM Na^+^) medium or hypertonic (300 mM Na^+^) medium for the appropriate time points. Quantified partition coefficients of DCP1A and G3BP are represented in the far right panel. (B) Cells were mock treated with 1x PBS or treated with 0.5 mM SA for the appropriate time points. Quantified partition coefficients of DCP1A and G3BP are represented in the far right panel. (C) Cells were first treated with hypertonic medium (300 mM Na^+^) for the appropriate time points and then rescued with isotonic (150 mM Na^+^) medium for various durations. Bars with green and red outline depict data points from panel A. (D) Cells were first treated with 0.5 mM SA for the appropriate time points and then rescued with medium containing 1x PBS for various durations. Bars with green and red outline depict data points from panel B. n = 3, > 60 cells, ***p ≤ 0.0001, N.S. denotes non-significance by two-tailed, unpaired Student’s t-test.

Next, we tested whether the increased focus number could be rescued (Res) by first subjecting cells to stress and subsequently recovering them in regular, isotonic growth medium (Figures 1C, 1D, S1E and S1F). We observed that hypertonicity induced DCP1A foci rapidly disappeared, within 2 min, irrespective of the duration of the stress (Figure 1C). While a significant fraction of the G3BP and polyA RNA foci also rapidly disappeared (within 2 min), the kinetics of complete recovery to the baseline (i.e., pre-treatment) focus number differed from those of DCP1A (Figures 1C and S1E). By comparison, DCP1A, G3BP, and polyA RNA foci induced by SA stress did not disappear even after 60 min of rescue (Figures 1D and S1F). These data suggest that DCP1A and G3BP/polyA RNA foci show differences in the kinetics of both assembly and disassembly, and that the rapid phase separation of DCP1A in response to altered tonicity is distinct from SG formation.

### Hypertonicity rapidly induces the formation of immobile DCP1A condensates in live cells

Since fixed-cell experiments revealed that the rapid phase separation of DCP1A condensates was distinct from SG formation over minutes and hours, we decided to probe the sub-cellular dynamics at greater temporal resolution. To this end, we subjected the previously developed UGD cell line (a U2OS cell line that stably expresses GFP-DCP1ADCP1A; (Pitchiaya et al., 2019) to a systematic set of hypertonic conditions. We chose this cell line because it contains a similar number of DCP1A foci as the parental U2OS cells, and each of these foci compositionally resembles endogenous PBs (Pitchiaya et al., 2019). As a control, we first confirmed in transiently transfected U2OS cells that DCP1A rapidly and reversibly forms “condensates” (Banani et al., 2017) irrespective of the fluorescence tag to which it is fused (GFP, mCherry, Halo, or CLIP; Figure S2A). We noted that the condensation and rescue of SNAP tagged DCP1A were distinct from the other tags (Figure S2A), raising the possibility that the nature of tagging might interfere with phase separation. Next, live cell imaging of UGD cells subjected to a cycle of isotonic conditions, brief hypertonic stress, and isotonic rescue (Movie S1) recapitulated the rapid and reversible nature of DCP1A phase separation (Figure 1). Furthermore, imaging of UGD cells at various levels of tonicity (150 mM to 450 mM Na^+^) showed that the number of GFP-DCP1A condensates per cell rapidly and monotonically increases with the salt concentration (Figure 2A); however, the mobility of the condensates, as measured by their diffusion constants, decreases. Within the time frame of treatment, typically 1-3 min, the cells remained viable across all concentrations of Na^+^ and survived 225 mM Na^+^ for up to 24 h (Figures 2B and S2B). Considering that many cellular processes depend on Mg^2+^ and Ca^2+^, we next examined whether DCP1A condensation was affected by increased concentration of these divalent metal ions in the growth medium. Both Mg^2+^ and Ca^2+^ induced GFP-DCP1A condensation only upon 100-fold increase over isotonic concentrations (Figures 2C and 2D), which corresponded to a significant increase in osmolarity to near double the osmolarity of isotonic growth medium (∼600 mOsm/L). These data suggest that DCP1A condensation occurs independently of the type of ion.

**Figure 2.**
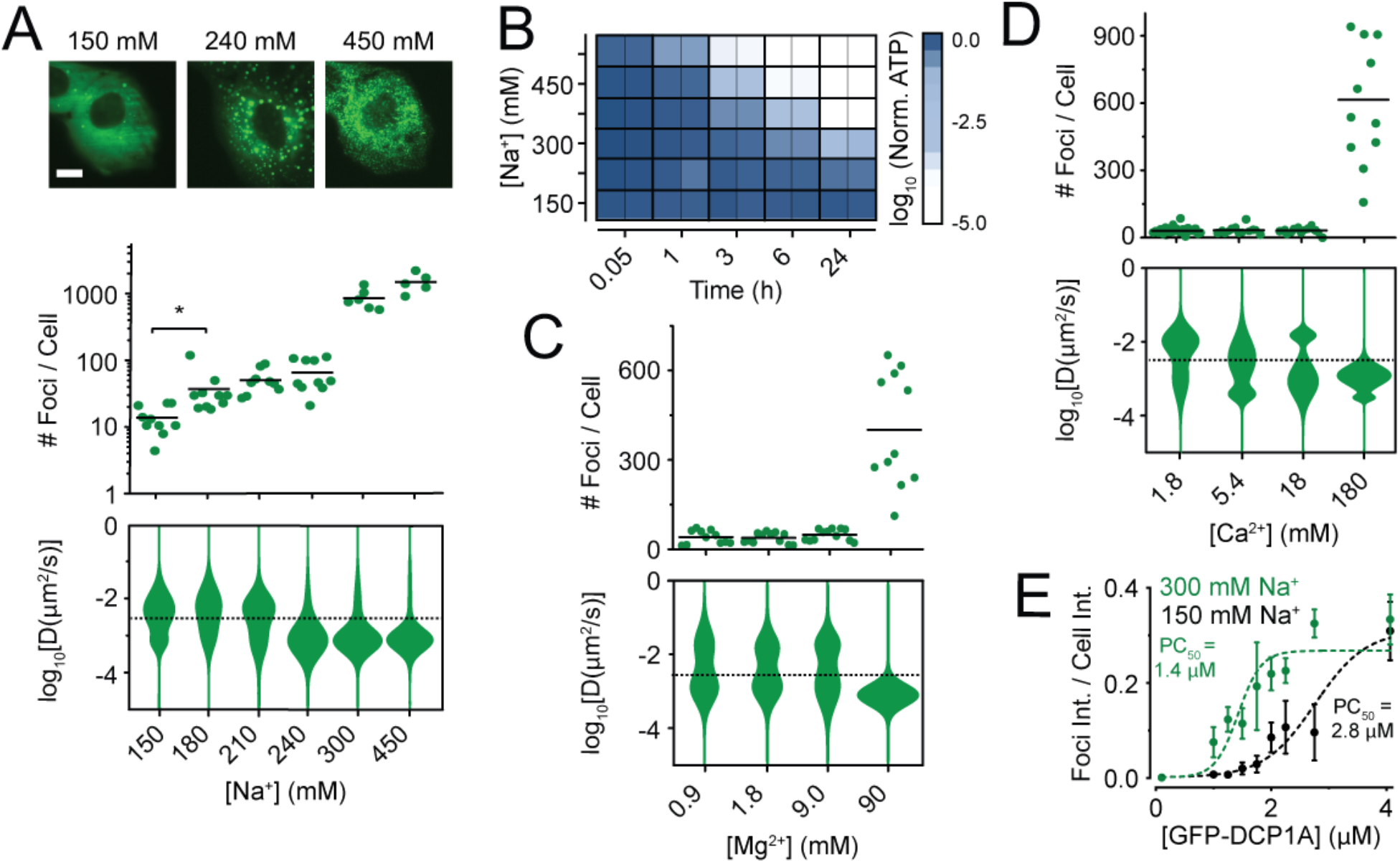
Physicochemical and phenotypic characterization of DCP1A phase separation during hypertonic stress. (A) Representative pseudocolored images of UGD cells (GFP, green) treated with growth medium containing various concentrations of Na+ (top), scatter plot of the number of foci per cell (middle), violin plots of diffusion constants associated with DCP1A foci (bottom). n = 2, > 5 cells per sample, *p ≤ 0.01 by two-tailed, unpaired Student’s t-test. (B) Representative image of a 96-well plate probed for cell viability by cell-titer glo assay across various Na^+^ concentrations and multiple time points. n = 3, with technical replicates for each n. (C and D) Scatter plot of the number of foci per cell (top) and violin plots of diffusion constants associated with DCP1A foci (bottom) within UGD cells treated with growth medium containing various levels if Mg^2+^ (C) or Ca^2+^ (D). n = 3, > 5 cells per sample. The dotted line in the diffusion plots empirically demarcates high- and low-mobility fractions. (E) Plot of partition coefficient against cellular concentration of DCP1A in samples treated with 150 mM Na^+^ (light green) or 300 mM Na^+^ (dark green). Data points were fitted with a dose response curve. PC50 = half maximal partition coefficient.

Biomolecular condensation generally has been found to scale with protein concentration (Banani et al., 2016; Sanders et al., 2020). To test whether hypertonicity-induced foci formation follows this paradigm, we measured the partition coefficient of GFP-DCP1A across a wide range of intracellular DCP1A expression levels. To this end, we calculated the partition coefficient (PC) as the fraction of total fluorescence assembled into discernable DCP1A foci and found that it scales linearly with the foci number (Figure S2C). We exploited this observation to show that the intracellular concentration leading to half-maximal partitioning was significantly lower in hypertonic conditions (PC50 = 1.4 µM, Figure 2E) than in isotonic controls (PC50 = 2.8 µM), supporting the notion that the extent of foci formation scales with protein concentration (Figure 2E). We then asked whether differences in their cellular concentration underlies differences in hypertonicity-induced condensation of DCP1A and G3BP. To this end, we compared GFP localization across U2-OS cells that stably expressed similar levels of either GFP-DCP1A (in UGD cells) or GFP-G3BP (UGG cells, Figures S2D and S2E). We found that, upon the characteristic short-term (2 min) hypertonic shock, GFP-DCP1A has a much higher foci forming potential (25-fold higher foci number and 30-fold higher partition coefficient) than GFP-G3BP. In contrast, foci formation in mock treated, short-term and prolonged SA treated controls (Figures S2D and S2E) were less prominent (except for GFP-G3BP upon prolonged SA treatment, as expected) and similar to those observed in the IF assays of Figures 1A, 1B, S1A, and S1C. These data are consistent with the notions that hypertonicity-induced DCP1A foci are not canonical SGs and that the differences observed between DCP1A and G3BP are due to specific sequence or structure motifs rather than their cellular abundance. Moreover, the rapid change in foci number generally occurs without a concomitant change in the total GFP fluorescence of the cell (Figure S2E), indicating that the GFP-DCP1A condensation is a direct response of the existing cellular protein to osmotic perturbation rather than an indirect response of protein expression.

### DCP1A phase separation is modulated by osmotic cell volume change

To distinguish between the possibilities that DCP1A condensation is a result of an increase in either specifically ionic or general osmotic concentration, we examined the sub-cellular effects of two non-ionic osmolytes, sucrose and sorbitol. Subjecting UGD cells to 300 mOsm/L of either of these osmolytes supplemented to regular growth medium again resulted in the formation of DCP1A condensates of lower mobility; however, cells recovered quickly when reversing to isosmotic medium (Figures 3A and S3A). These observations strongly suggest that DCP1A condensates form in response to osmotic shock rather than changes in ionic strength only.

**Figure 3.**
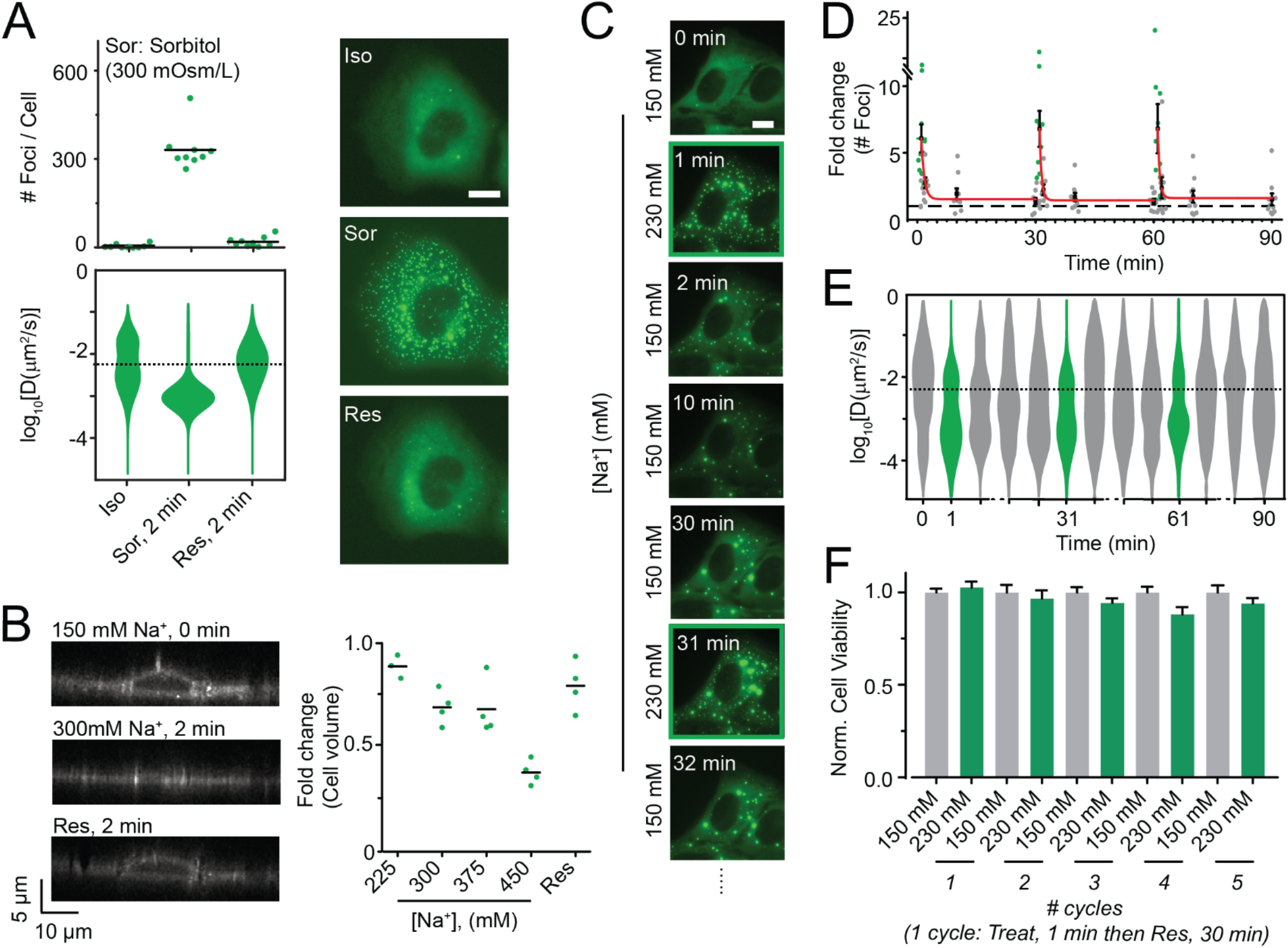
Hyperosmotic compression mediates DCP1A phase separation. (A) Scatter plot of the number of foci per cell (top), violin plots of diffusion constants associated with DCP1A foci (bottom) and representative pseudocolored images of UGD cells (GFP, green) treated with isosmotic (Iso) growth medium, hyperosmotic growth medium containing the non-ionic osmolyte Sorbitol (Sor), or rescued (Res) with isosmotic medium after Sorbitol treatment. n = 2, > 5 cells per sample. Scale bar, 10 µm. (B) Representative y-z projection of UGD cells (gray-scale) from 3-D imaging assay wherein the cell were treated with isotonic (150 mM Na^+^) medium, hypertonic (300 mM Na^+^) medium or rescued with isotonic medium after hypertonic treatment. n = 1, 4 cells per sample. Scale bar, 10 µm. Scatter plot of the fold change in cell volume, as normalized to the cell volume in isotonic conditions, is shown. (C) Representative pseudocolored images of a UGD cell (GFP, green) that was cyclically treated with isotonic (150 mM Na^+^) or hypertonic (300 mM Na^+^) medium. Scale bar, 10 µm. (D) Scatter plot of the fold change in foci number, as normalized to foci number in isotonic samples, associated with assay represented in C. Red line depicts exponential fit. n = 2, > 5 cells per sample. (E) Violin plots of diffusion constants associated with DCP1A foci, associated with assay represented in C. n = 2, > 5 cells per sample. The dotted line in the diffusion plots empirically demarcates high- and low-mobility fractions. (F) Bar plots of cell viability (by CellTiter-Glo assay), as normalized to isotonic samples, associated with assay represented in C. n = 3, with 3 technical replicates for each n.

Since hyperosmolarity is a state of increased extracellular osmotic pressure and causes cellular volume reduction by compensatory exosmosis (i.e., water loss), we hypothesized that DCP1A foci formation is the result of osmotic cellular compression causing an increase in intracellular molecular crowding (Guo et al., 2017; Miermont et al., 2013). To test this hypothesis, we estimated volume changes of UGD cells in hyperosmotic medium using DiI staining (Sukenik et al., 2018) and 3-dimensional (3-D) live-cell and fixed-cell imaging (Figures 3B, S3B and S3C). We found that cell height, as a proxy for cell volume, rapidly (within ∼1 min) and monotonically decreased over the increasing range of tested osmotic conditions in live-cell imaging conditions with a concomitant increase in the number of DCP1A foci. This hypertonicity-induced rapid reduction in cell volume was also observed by fixed-cell 3-D imaging and flow cytometry (Figures S3C and S3D), although with reduced sensitivity, owing to fixation-induced cell shrinkage (Ross, 1953; Su et al., 2014). Since osmotic volume change has previously been reported to be reversible in response to prolonged osmotic stress (Burg et al., 1997; Hoffmann et al., 2009), we next asked whether regulatory volume increase occurs on the timescales at which we observe formation and persistence of GFP-DCP1A condensation. However, cell height reduction in fixed cells showed little change upon prolonged hyperosmolarity, going from ∼20% (10 min) to ∼25% (120 min) relative to isosmotic controls (Figure S3E), suggesting that volume recovery does not play a significant role in our system (Burg et al., 1997; Hoffmann et al., 2009). Moreover, the effect of hyperosmotic volume compression on DCP1A foci formation was independent of cell lineage (Figure S3F). Together, our data support a direct, universal link between molecular crowding and GFP-DCP1A condensation.

Since DCP1A exhibits rapid and reversible condensation dependent on the degree of osmotic cell volume change, and since mammalian cells repeatedly experience such osmotic perturbations, we examined the response of UGD cells to cycling osmotic volume change (Figures 3C and 3D). UGD cells were treated for 1 min with hypertonic medium and allowed to recover for 30 min, and this treatment regimen was repeated. Quantification of the number and diffusion constants of foci across the treatment regimen showed that the time-scales of DCP1A focus assembly (∼10 s) and disassembly (∼100 s), as well as changes in focus mobility, were highly similar across all cycles and occurred with minimal impact on cell viability (Figures 3C-3F). We henceforth refer to this phenomenon of cytosolic DCP1A condensation as intracellular hyperosmotic phase separation (HOPS) and posit that it is a cellular adaptation to osmolarity-induced changes in molecular crowding.

### HOPS of DCP1A depends on its trimerization domain and post-translational modification status

Macromolecular phase separation is widely thought to be driven by multivalent protein-protein and protein-nucleic acid interactions mediated by specific side chain interactions and structures (Guo and Shorter, 2015). To investigate the underlying molecular basis of DCP1A condensation, we first tested the dependence of HOPS on different DCP1A domains. While DCP1A does not contain any annotated nucleic acid binding domains, it contains two prominent protein interaction domains, an N-terminal EVH1 domain that interacts with the mRNA decapping protein DCP2, a C-terminal trimerization domain that interacts with EDC3/4, and a scaffolding protein of the decapping complex (Aizer et al., 2013). GFP-tagged truncation constructs of DCP1A’s N-terminal domain (NTD) or C-terminal domain (CTD) were transiently transfected into U2OS cells. Upon exposing these cells to hyperosmotic shock, we observed that the CTD showed rapid and reversible condensation similar to the full-length protein. In contrast, a truncation mutant containing the NTD did not show detectable foci upon hyperosmotic shock (Figures 4A and 4B). As the CTD mediates both DCP1A trimerization and EDC4 interaction, we tested whether EDC4 is responsible for HOPS of DCP1A to narrow down the basis of condensation. Compared to a scrambled (Scr) silencing RNA (siRNA) control, knockdown of EDC4 by siEDC4 resulted in reduced expression (∼2-fold) of GFP-DCP1A (Figures S4A and S4B) and larger GFP-DCP1A foci under isotonic conditions (Figures 4C and 4D), but did not prevent HOPS of DCP1A. In fact, the slight reduction in HOPS of DCP1A is consistent with the ∼2-fold reduced cytosolic availability of DCP1A via reduced expression and enhanced localization within large foci. These data strongly suggest that DCP1A homo-trimerization is a major driver of its HOPS.

**Figure 4.**
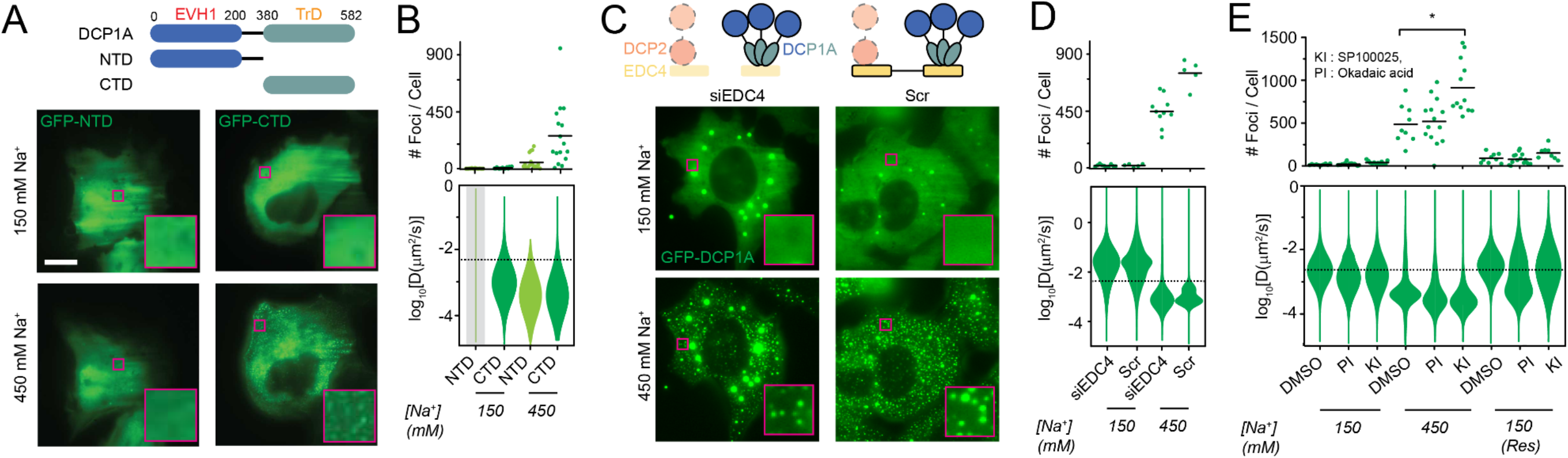
HOPS of DCP1A is dependent on its trimerization domain and modulated by PTMs, but not its interaction with EDC4. (A) Schematic of full length DCP1A, NTD, or CTD constructs (top, not to scale). EVH1 domain, trimerization domain, and the amino acid numbers are marked. Representative pseudocolored images of U2OS cells (GFP, green) transfected with GFP-NTD or GFP-CTD that were treated with isotonic (150 mM Na^+^) or hypertonic (300 mM Na^+^) medium (bottom). Scale bar, 10 µm. (B) Scatter plot of the number of foci per cell (top) and violin plots of diffusion constants associated with DCP1A foci (bottom) imaged in panel A. n = 3, > 5 cells per sample. (C) Schematic of DCP1A, DCP2 and EDC4 in the decapping complex (top, not to scale) in siEDC4 or Scr treatment conditions. Representative pseudocolored images of siEDC4 or Scr siRNA treated UGD cells (GFP, green) treated with isotonic (150 mM Na^+^) or hypertonic (300 mM Na^+^) medium (bottom). Scaled as in panel A. (D) Scatter plot of the number of foci per cell (top) and violin plots of DCP1A diffusion constants (bottom), associated with assay represented in C. n = 3, > 5 cells per sample. (E) Scatter plot of the number of foci per cell (top) and violin plots of DCP1A diffusion constants (bottom) within UGD cells that were pre-treated treated with DMSO, KI, or PI, and imaged in isotonic (150 mM Na^+^) medium, hypertonic (300 mM Na^+^) medium, or rescued (Res) with isotonic medium after hypertonic treatment. n = 3, > 5 cells per sample. The dotted line in the diffusion plots empirically demarcates high- and low-mobility fractions.

Previous reports have suggested that PB formation can be modulated by post-translational modification (PTM), as accompanying cell cycle progression (Aizer et al., 2008). We reasoned that if change in phosphorylation status would influence PB assembly and disassembly during the cell cycle, such modifications should also modify the protein’s response to changes in molecular crowding. We stimulated global changes in phosphorylation using either a general phosphatase inhibitor (PI), okadaic acid, or the c-Jun N-terminal kinase inhibitor (KI) SP600125 on UGD cells (Aizer et al.), which we found to modulate DCP1A phosphorylation levels (Figure S4C). While the PI did not significantly alter HOPS of GFP-DCP1A compared to a DMSO treated control, the KI significantly increased the number of newly formed immobile condensates (Figure 4E). Additionally, PI treatment mediated a significant reduction in the mobility of DCP1A condensates even after rescuing the cells with isotonic medium (Figure 4E). Together, these observations suggest that PTMs can modulate HOPS of proteins, likely by altering surface charges that protein-protein interactions depend on.

### Multimeric proteins with a valency of at least 2 generally exhibit HOPS

Considering that the minimally required structural determinant of HOPS of DCP1A is only its trimerization domain (Figure 4), we reasoned that other self-interacting proteins with multimerization domains might also exhibit HOPS. To test this hypothesis, we performed an unbiased high-throughput IF analysis of 104 endogenous proteins and 4 nucleic acids-based targets (m6A, RNA-DNA hybrid poly-ADP-ribose and rRNA) in U2OS cells subjected to short-term osmotic stress (Figures 5A, 5B and S5A, xref Table S1). Since antibodies may exhibit cross-reactivity and impaired access to some proteins in osmotically compressed cells, we complemented high-throughput IF analysis by imaging osmotically perturbed U2OS cells transiently transfected with GFP-tagged proteins (Figures 5C, S5B and S5C and Methods). A combined analysis of both assays showed that proteins annotated as multimers or homooligomers (by gene ontology) that were trimers or larger had a higher propensity to exhibit HOPS (Figure 5C-5E). For instance, monomeric proteins (e.g., GFP), dimeric proteins (e.g., TP53, AKT), and several proteins without annotated multimerization domains (e.g., PARP13) did not exhibit HOPS (65 out of 75 targets, Figures 5C-5E). By contrast, a significant fraction (18 out of 29 annotated homo-oligomers) of multimeric proteins with a self-interaction valence of ≥2 (i.e. trimers and other higher-order multimers, including LCD bearing proteins such as DCP1A, HSF1, PKM2, PAICS, FUS, and TDP43), as well as several proteins with no known multimerization domain (e.g., ERp72) exhibited HOPS (Figures 5C-5E and S5). In some cases, we observed the disappearance of foci upon hyperosmotic shock, which rapidly reformed upon isotonic rescue (e.g., CDK12; Figures 5B and S5A). These observations support the hypothesis that multimeric proteins with a self-interaction valency of 2 or more generally undergo HOPS. Overall, our data suggest that the sub-cellular distribution of a significant fraction of the cellular proteome, 16% of which is annotated to be self-interacting (Perez-Bercoff et al., 2010), appears to be altered by osmotic compression. Our findings thus support a widespread and pervasive impact of HOPS on subcellular organization. We then tested whether hetero-multimers exhibit HOPS, by using live cell imaging of cells co-expressing Halo-tagged DCP1A and GFP-tagged interactors of DCP1A. While overexpression of GFP-AGO2, which co-localizes with DCP1A at PBs, and GFP-DCP2, which directly interacts with DCP1A, by themselves failed to undergo HOPS, high co-expression with Halo-DCP1A induced GFP-AGO2 and GFP-DCP2 condensates that colocalized with DCP1A HOPS condensates (Figure S6). Notably, using GFP-DCP1A and Halo-DCP1A co-expression as a positive control showed significantly higher HOPS propensity, as expected for DCP1A, whereas GFP by itself failed to undergo condensation under any condition (Figure S6). We conclude that trimeric DCP1A is sufficient to drive the formation of these condensates, but that HOPS can also be induced via hetero-multimerization dependent on the extent of interaction and concentration of the interactors. At endogenous concentrations of PB constituents, DCP1A HOPS condensates may represent primarily DCP1A protein (Figure S1B).

**Figure 5.**
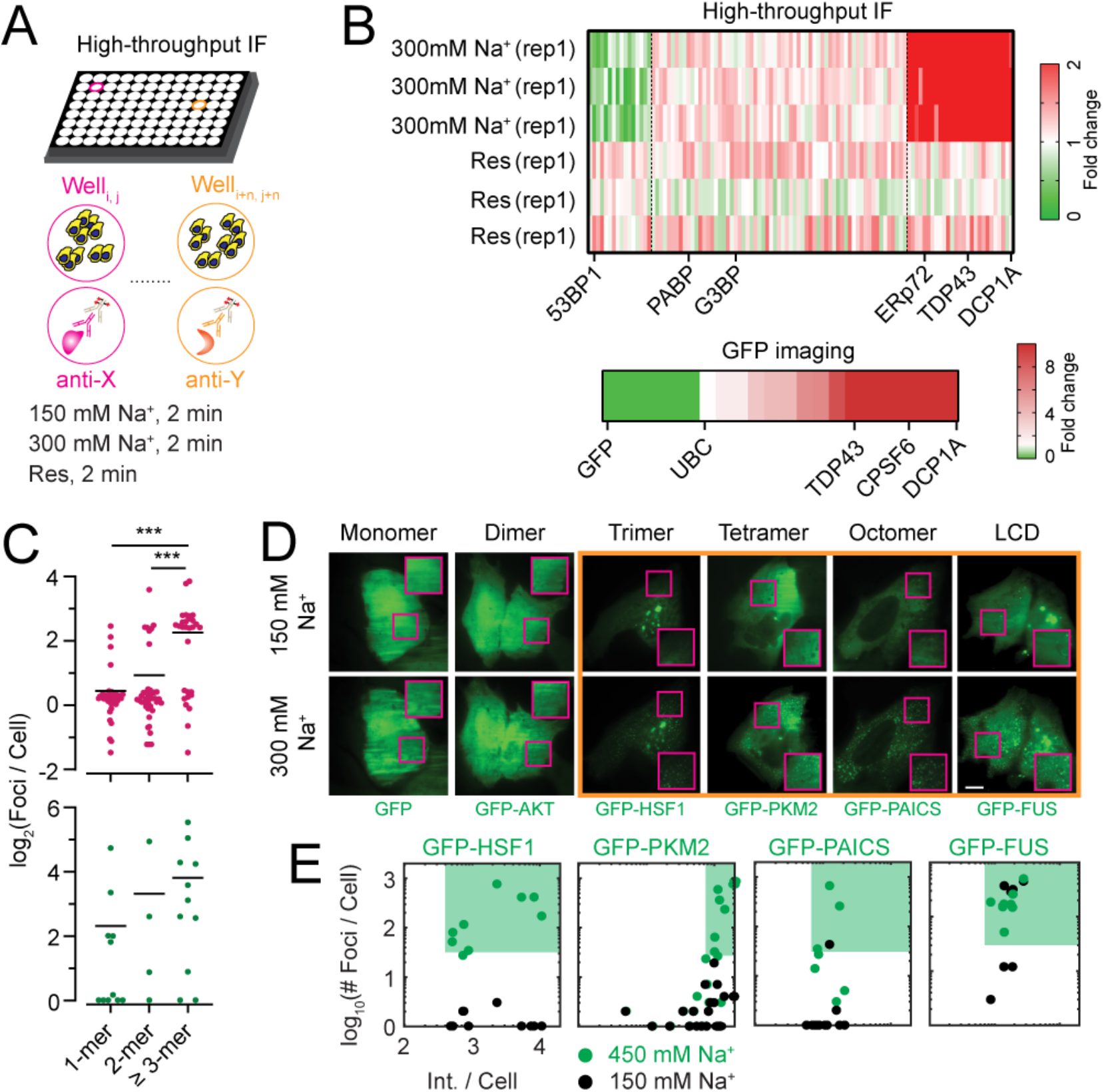
High-throughput IF and GFP imaging show that several multimeric proteins of valency ≥2 generally exhibit HOPS. (A) Schematic of high throughput IF assay. (B) Heatmap representing the fold change in spot number of the 108 targets tested by high throughput IF, as normalized to isotonic conditions. “rep” denotes replicates. Heatmap representing the fold change in spot number of 22 protein targets tested by GFP imaging, normalized to isotonic conditions are shown below. (C) Scatter plot of the average number of foci per cell as a function of known protein multimerization ability. Each dot represents an individual target in the high-throughput IF of GFP imaging assay. n = 3, ***p ≤ 0.0001 by two-tailed, unpaired Student’s t-test. (D) Representative pseudocolored images of U2OS cells (GFP, green) transfected with the appropriate GFP-tagged construct and treated with isotonic (150 mM Na^+^) medium or hypertonic (300 mM Na^+^) medium. Scale bar, 10 µm. Inset depicts a zoomed-in area corresponding to a 15 x 15 µm^2^ magenta box. The orange box encloses proteins that exhibit HOPS. (E) Scatter plot of the number of foci per cell against the area-normalized cell intensity for each protein is shown. n = 2, > 5 cells per sample. The green contour depicts the HOPS regime.

### HOPS of CPSF6 is correlated with hyperosmolarity-induced impairment of transcription termination

One of the HOPS positive hits in our High-throughput IF and GFP imaging was CPSF6 (Figures 5B and 6A), a structural component of the cleavage and polyadenylation specific factors (CPSFs) containing complex (Elkon et al., 2013), which in turn mediates 3’end formation of transcripts. Prior reports have shown that impaired function of CPSFs during hyperosmotic stress is correlated with transcription termination defects in certain cellular lineages, leading to continued transcription of regions downstream of annotated genes (Vilborg et al., 2015). We posited that the sequestration of CPSF6 from its site of action (transcription sites) by HOPS mediates transcription termination defects under hyperosmotic stress. To test this hypothesis, we assessed how hyperosmolarity affected the nascent transcriptome, which was expected to be highly sensitive to termination defects, using nascent state RNA sequencing of transcripts by 5-bromouridine metabolic labeling and sequencing (Bru-seq) and BruChase-seq after 30 min of hyperosmotic stress (Paulsen et al., 2014). We found that, indeed, the read density of sequences downstream of transcription end sites (TES) was significantly higher in the hypertonic samples than under isotonic conditions (Figures 6B, 6C and S7A). Performing steady-state RNA-seq of UGD cells subjected to prolonged (4 h) osmotic perturbations revealed that hyperosmotic stress also had a pervasive long-term effect that, strikingly, was reversed upon rescuing cells from hypertonic shock with isotonic medium (Figures S7B and S7C). Chromatin-immunoprecipitation followed by sequencing (ChIP-seq) further showed that elongating RNA Pol II was enriched downstream of canonical TES specifically in the hyperosmotic samples, consistent with transcription proceeding beyond the TES (Figure S7D). In addition, ChIP-seq analysis of CPSF proteins CPSF-1, CPSF6 and CPSF73 (Figures 6D, S6E and S6F) showed that the localization of CPSF complexes at TES was significantly reduced under hyperosmotic conditions. Taken together, a model emerges by which rapid formation of HOPS condensates of CPSF6 sequesters CPSF activity away from chromatin, leading to the functional impairment of cleavage and polyadenylation at the TES of a subset of transcripts upon hyperosmotic stress (Figure 6D).

**Figure 6.**
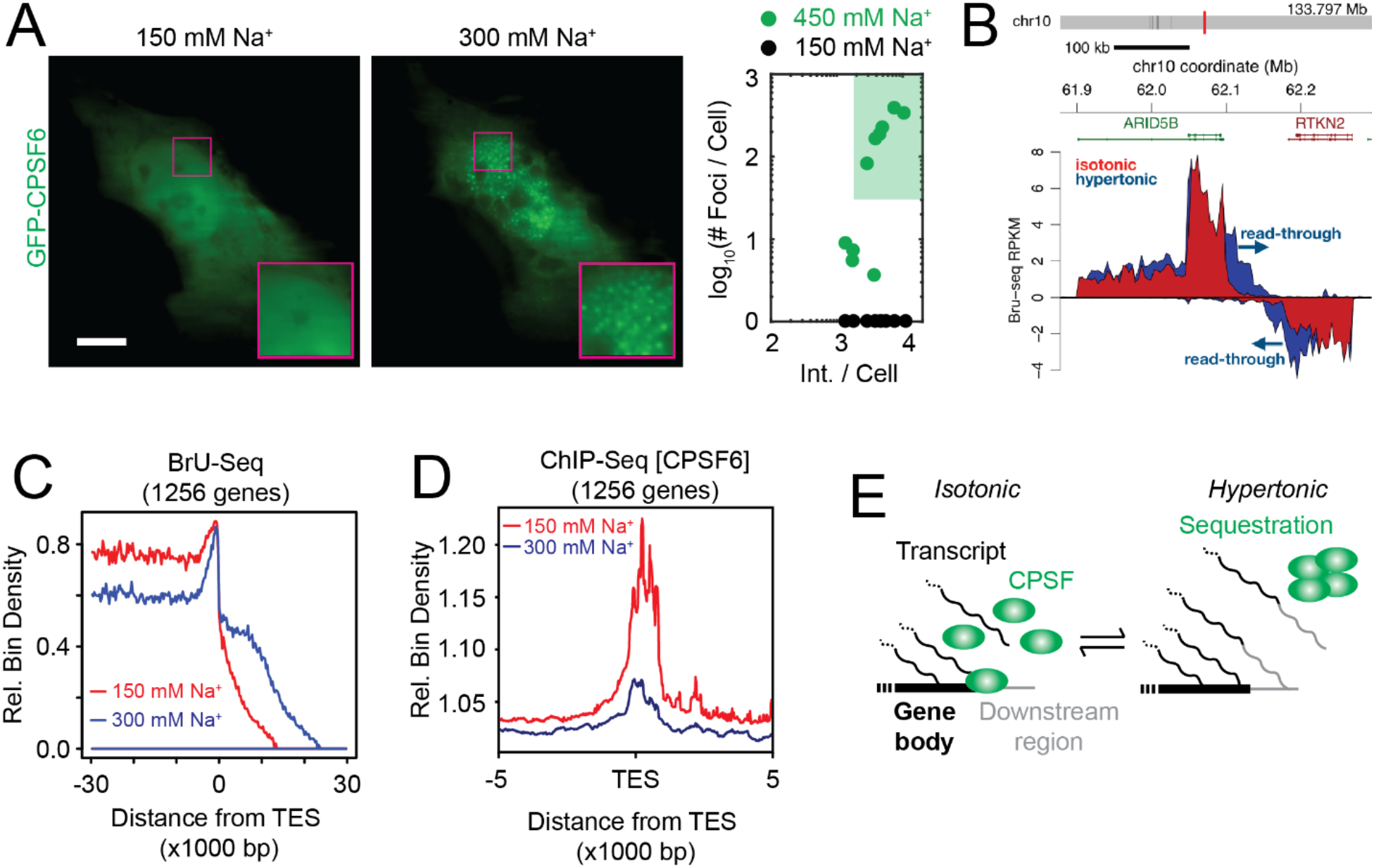
HOPS of CPSF6 is correlated with impaired transcription termination. (**A**) Representative pseudocolored images of a U2OS cell transfected with GFP-CPSF6 (green) incubated with isotonic (150 mM Na^+^, red) medium and then treated with hypertonic (300 mM Na^+^, blue) medium for 1 min. Scale bar, 10 µm. (B) Bru-seq tracks across the ARID5B and RTKN2 genes showing transcriptional read-through of the TES. (C) Aggregate nascent RNA Bru-Seq enrichment profile across TESs. Relative bin density of ∼1256 genes expressed >0.5 RPKM and >30 kb long showing an ∼10 kb average extension of reads past the TES following exposure to hypertonic conditions for 30 min. Samples were prepared from cells treated with isotonic (150 mM Na^+^, red) or hypertonic (300 mM Na^+^, blue) medium for 30 min. (D) Aggregated ChIP-seq peaks of CPSF6 around the TES under hypertonic (300 mM Na^+^, blue) and isotonic conditions (300 mM Na^+^, red). (E) Schematic model of transcription termination defect induced by HOPS of CPSFs.

## DISCUSSION

In this study, we report a multiscale (i.e., cellular, subcellular, and molecular) characterization of a seemingly widespread intracellular phase separation phenomenon in response to hyperosmotic stress, here termed HOPS (Figure 7). We find that a significant fraction of the multimeric proteome undergoes rapid and reversible intracellular redistribution into phase-separated condensates during osmotic cell volume change. Empirically, proteins with a self-interaction valency of ≥2 exhibit HOPS in response to changes in cell volume, and these changes are in turn intricately linked with altered hydration and molecular crowding during hyperosmotic stress.

**Figure 7.**
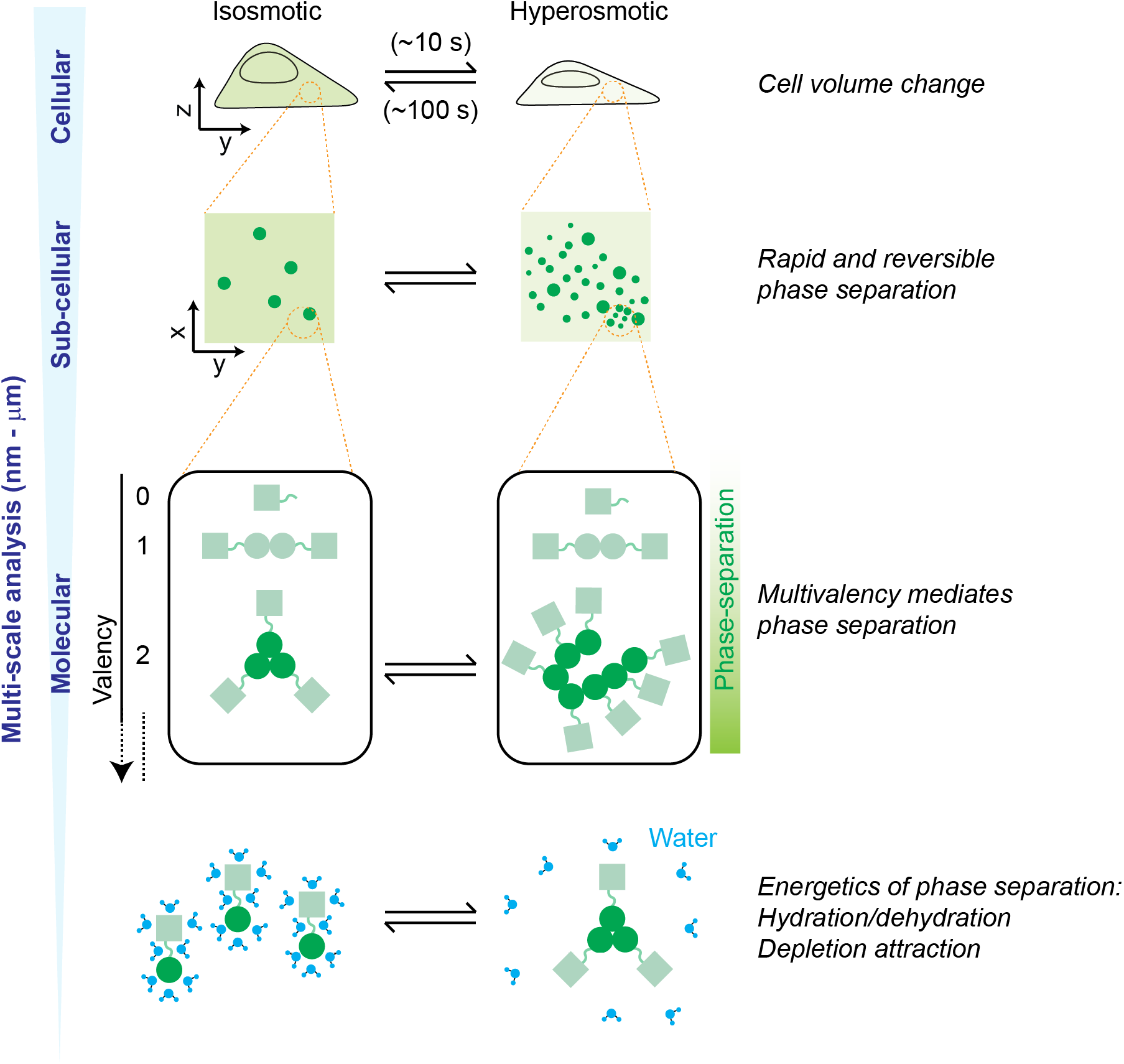
Model of the multiscale features of HOPS. Our multi-scale analysis has shown that HOPS of multimeric proteins is mediated by the concerted changes in cell volume, macromolecular crowding, and hydration.

### Exosmosis leads to protein concentration increase, molecular dehydration, and HOPS

Intracellular water expelled upon hyperosmotic compression (i.e. exosmosis) is thought to originate from both “free” water molecules that constitute the bulk of the cell and water molecules bound to cellular solutes and involved in macromolecular solvation (Ball, 2017). On the one hand, the loss of free water upon exosmosis leads to cell volume loss and a concomitant increase in cellular concentration that will shift the monomer-multimer equilibrium of a protein towards multimerization, which may be facilitated by depletion attraction (Marenduzzo et al., 2006). On the other hand, the loss of bound water will result in decreased protein hydration, which may lead to protein precipitation by increasing the surface exposure of hydrophobic regions (Muschol and Rosenberger, 1997). Rehydration rapidly replenishes both types of water molecules to shift the monomer-multimer equilibrium back towards the solvated monomer, dissolving the condensate.

It is thought that hydrophobic patches found in homo-multimeric proteins can spontaneously interact upon hydration loss or “dewetting” (Jensen et al., 2003; Liu et al., 2005), whereas the phase separation driven by LCDs and RNAs is posited to involve larger interaction networks (Wang et al., 2018). The high speed and high reversibility of HOPS of multimeric proteins is consistent with the former mechanism wherein hydrophobic patches of homo-oligomeric proteins promote association into condensates (Krainer et al., 2020) (Figure 7). The cost in translational entropy of individual diffusing molecules upon condensation into such large, slowly diffusing complexes is expected to be compensated by the enthalpic gain of hydrophobic patch association.

## The features and functional consequences of widespread intracellular HOPS

We observe that increasing the intracellular crowding ∼2-fold (based on the up to ∼2-fold change in cell height) leads to the formation of a large number of DCP1A condensates with greatly reduced mobility; further, cellular volume recovery readily reverses both the condensation and decreased mobility (Figures 2 and 3). Additionally, we find that the cellular concentration of the protein monomer affects the size and number of condensates (Figure S4D). The latter observation implies that, under low protein concentration conditions, our ability to identify proteins undergoing HOPS may be limited by our fluorescence microscope’s resolution. A conservative estimate, based on cytoplasmic redistribution of GFP signal into hyperosmotic condensates (Figures S2D and S2E), suggests that we can detect 10-mers and any higher-order condensates. This level of sensitivity has allowed us to use IF to curate a high-confidence list of endogenous proteins that do and do not undergo HOPS (Figure 5). Thus, we can define the protein features that govern HOPS, primarily the requirement for a homo-multimerization domain of valency ≥2 that is common among cellular proteins.

It is becoming increasingly clear that excluded volume effects mediated by molecular crowding affect macromolecular structure, protein stability, enzyme activity and nucleo-cytoplasmic organization (Daher et al., 2018; Delarue et al., 2018; Hancock, 2004; Minton, 2001; Sukenik et al., 2018). Previous work has noted the potential for phase separation to dynamically buffer the intracellular protein concentration (Alberti et al., 2019). More directly, we find that the structural pre-mRNA cleavage and polyadenylation factor CPSF6 (Elkon et al., 2013) undergoes nuclear HOPS, which we observe to be correlated with transcriptome-wide functional impairment of transcription termination (Figure 6).

### HOPS may serve as a rapid cellular sensor of volume compression

The rapid time scales of hyperosmotic cell volume compression, volume recovery under isotonicity, and cell viability after multiple osmotic cycles (Figures 2 and 3) that we observe concur with prior reports on cell volume changes (Guo et al.; Hersen et al.; Miermont et al.). Our data, which indicate that even a 20% reduction in cell volume by osmotic compression can mediate HOPS, reinforce evidence of the high sensitivity of the multimeric proteome towards volume changes. Our findings are consistent with the notion that the eukaryotic proteome is delicately balanced near the threshold of phase separation (Walter and Brooks, 1995; Wilson, 1899). In fact, it stands to reason that the interaction energies and concentrations of homo-multimeric proteins may have evolved to facilitate rapid crossing of their individual phase separation thresholds if, and only if, cellular conditions demand.

Notably, our HOPS-associated cell volume changes are comparable to the rapid volume changes – also a result of exosmosis – occurring during cell adhesion and migration through confined spaces (Guo et al., 2017; Watkins and Sontheimer, 2011), as well as those associated with the cell cycle (Tzur et al., 2009). Incidentally, homeostatic processes that may be expected to suppress phase separation, such as PTMs and allosteric effects by metabolically compatible osmolytes, operate over the time scales (minutes to hours) of the cell cycle. Consistent with this expectation, we find that the loss of phosphorylation enhances the extent of HOPS (Figure 4).

Perhaps the most striking aspect of HOPS is its rapid onset, which is faster than the rate of assembly of stress-response associated granules (Wheeler and Hyman, 2018). This feature is similar to recent reports of rapid nuclear condensation of DEAD-box RNA helicase DDX4 in response to environmental stress (Nott et al., 2015), of transcriptional co-activator YAP in response to hyperosmotic stress (Cai et al., 2019) and may be homologous to a similar phenomenon reported in yeast (Alexandrov et al., 2019). Notably, prolonged exposure to hyperosmotic conditions, similar to other environmental stressors, triggers the ISR and subsequent assembly of SGs, often localized adjacently to pre-formed DCP1A condensates or PBs (Figure 1) (Kedersha et al., 2005). These observations support a model whereby homo-multimeric proteins serve as “first responders” of osmotic compression due to the sensitivity of protein-complex formation to changes in crowding (Sukenik et al., 2017). Such early volume sensors may be critical for suspending cellular biochemistry until an appropriate protective or corrective action has been initiated. This escalating response may be critical since osmotic changes in the environment are unpredictable and can rapidly fluctuate, yet have widespread implications in an array of physiological and disease contexts. For instance, cells in the renal medulla frequently and rapidly experience high salt concentrations resulting in up to four-times the osmolarity of serum during urine production (Lang et al., 1998). Extreme dehydration can lead to hypernatremia, a state of serum hyperosmolarity characterized by elevated Na^+^ levels exceeding 145 mM, and is associated with pervasive physiological dysfunction (Nilsson and Sunnerhagen, 2011). During such prolonged stress, initiation of the ISR may then lead to long-term adaptation. For instance, long-lasting condensates of the protein WNK1, which notably also is a homo-multimer, have been observed in viable kidneys of mice raised on high-K^+^ diets (Boyd-Shiwarski et al., 2018).

Both acute and prolonged HOPS are reversible (Figure 1) and can mediate widespread effects, including impairment of transcription termination (Figure 6) (Vilborg et al., 2015), YAP-programmed transcription initiation (Cai et al., 2018), inhibition of ribosomal translocation (Wu et al., 2019), modulation of RNA silencing (Pitchiaya et al., 2019) and altered degradation of ribosomal proteins (Yasuda et al., 2020), all of which potentially contribute towards cell survival. While other mechanisms may be also at play, protein sequestration away from the site of their function provides a straightforward biophysical explanation for many of these effects (Figure 6). In fact, such a mechanism may also explain the defects in transcription termination observed in cells exposed to prolonged heat shock (Cardiello et al., 2018), suggesting that protein sequestration might be a common mechanism across multiple stress responses. Additionally, the rapid formation of rigid condensates may aid in maintaining the structural integrity of cell by altering physical forces in the crowded interior (Quiroz et al., 2020). Future studies will help better understand the connection between MLO formation and protective cellular mechanisms heralded here.

## Supporting information

Supplementary Reference List

## ACKNOWLEDGEMENTS

APJ was supported by NIH T-32-GM007315. SP was supported by an AACR prostate cancer research fellowship and an NCI-SPORE Career Enhancement Award. A.M.C. is a NCI Outstanding Investigator, Howard Hughes Medical Institute Investigator, A. Alfred Taubman Scholar, and American Cancer Society Professor. This work was supported by NIH grant R01 GM122803 and a University of Michigan Comprehensive Cancer Center/Biointerfaces Institute research grant to N.G.W and NCI Prostate SPORE (P50 CA186786) to A.M.C. We thank X. Cao, F. Su and R. Wang for technical assistance with sequencing library preparation, and M. Denies for help with some initial experiments. We also acknowledge NSF MRI-ID grant DBI-0959823 (to N.G.W.) for seeding the Single Molecule Analysis in Real-Time (SMART) Center, whose Single Particle Tracker TIRFM equipment was used for much of this study with support from J.D. Hoff. Finally, we would like to thank N. Kedersha and Y. Shav-Tal as sources of several plasmids.

## AUTHOR CONTRIBUTIONS

S.P. conceived the study. A.P.J. performed all live-cell imaging experiments. S.P. and X.J. performed all fixed cell assays and phenotypic analyses. L.X. constructed several plasmids. A.P., P.B., K.B., M.C. and M.L. performed and analyzed the sequencing assays. S.P., A.P.J., A.M.C. and N.G.W. designed the study. S.P., A.P.J. and N.G.W. and A.M.C. wrote the manuscript, and all authors provided feedback on the manuscript.

## SUPPLEMENTAL INFORMATION

**Multivalent proteins rapidly and reversibly phase-separate upon osmotic cell volume change**

Ameya P. Jalihal, Sethuramasundaram Pitchiaya, Lanbo Xiao, Pushpinder Bawa, Xia Jiang, Karan Bedi, Abhijit Parolia, Marcin Cieslik, Mats Ljungman, Arul M. Chinnaiyan and Nils G. Walter

## SUPPLEMENTARY FIGURES AND LEGENDS

**Figure S1.**
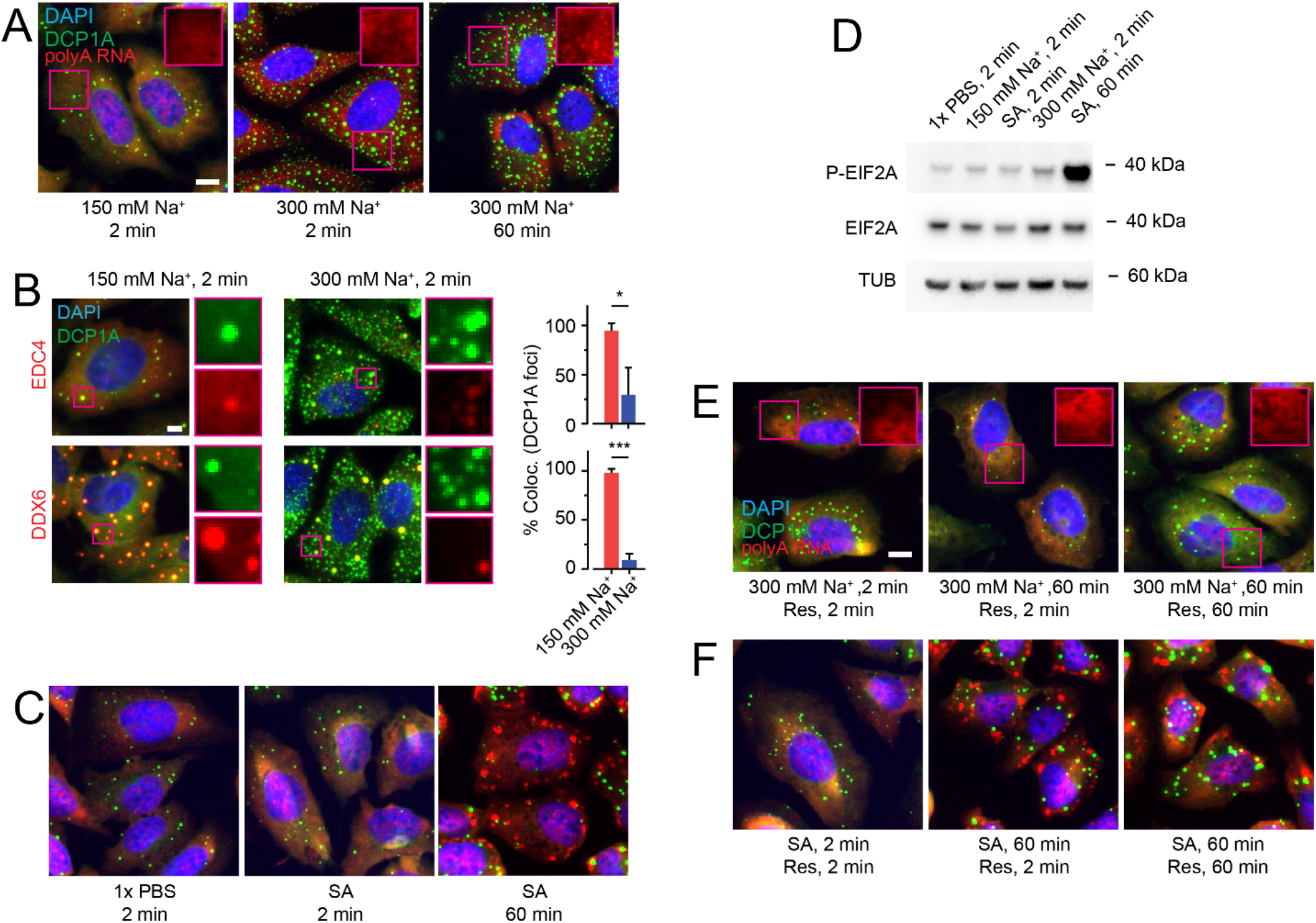
Hypertonicity-induced DCP1A foci are not canonical PBs or SGs. **Related to Figure 1.** (A) Representative pseudocolored, combined IF – RNA-FISH images of U2-OS cells stained for DAPI (blue), DCP1A (green) and polyA RNA (red) at various time points after isotonic (150 mM Na^+^) or hypertonic (300 mM Na^+^) medium addition. Scale bar, 10 µm. (B) Representative pseudocolored IF images of U2-OS cells stained for DAPI (blue), DCP1A (green), and EDC4 (top panels, red) or DDX6 (bottom panels, red) as indicated, treated with isotonic (150 mM Na^+^) or hypertonic (300 mM Na^+^) medium for 2 min. Scale bar, 10 µm. The plots represent the fraction of DCP1A foci colocalizing with EDC4 or DDX6 foci under each condition. n = 3, 50 cells, *p < 0.01, ***p < 0.0001, significance by two-tailed, unpaired Student’s t-test. (C) Representative pseudocolored, combined IF – RNA-FISH images of U2-OS cells stained for DAPI (blue), DCP1A (green) and polyA RNA (red), and either mock treated with 1x PBS or treated with 0.5 mM SA for the appropriate time points. (D) Western blot of phosphorylated EIF4E (P-EIF4E), EIF4E and tubulin upon various osmotic and SA perturbations. (E-F) Representative pseudocolored, combined IF – RNA-FISH images of U2-OS cells stained for DAPI (blue), DCP1A (green) and polyA RNA (red). Scale bar, 10 µm. (E) Cells were first treated with hypertonic (300 mM Na+) medium for the appropriate time points and then rescued with isotonic (150 mM Na+) medium for various durations. (F) Cells were first treated with 0.5 mM SA for the appropriate time points and then rescued with medium containing 1x PBS for various durations.

**Figure S2.**
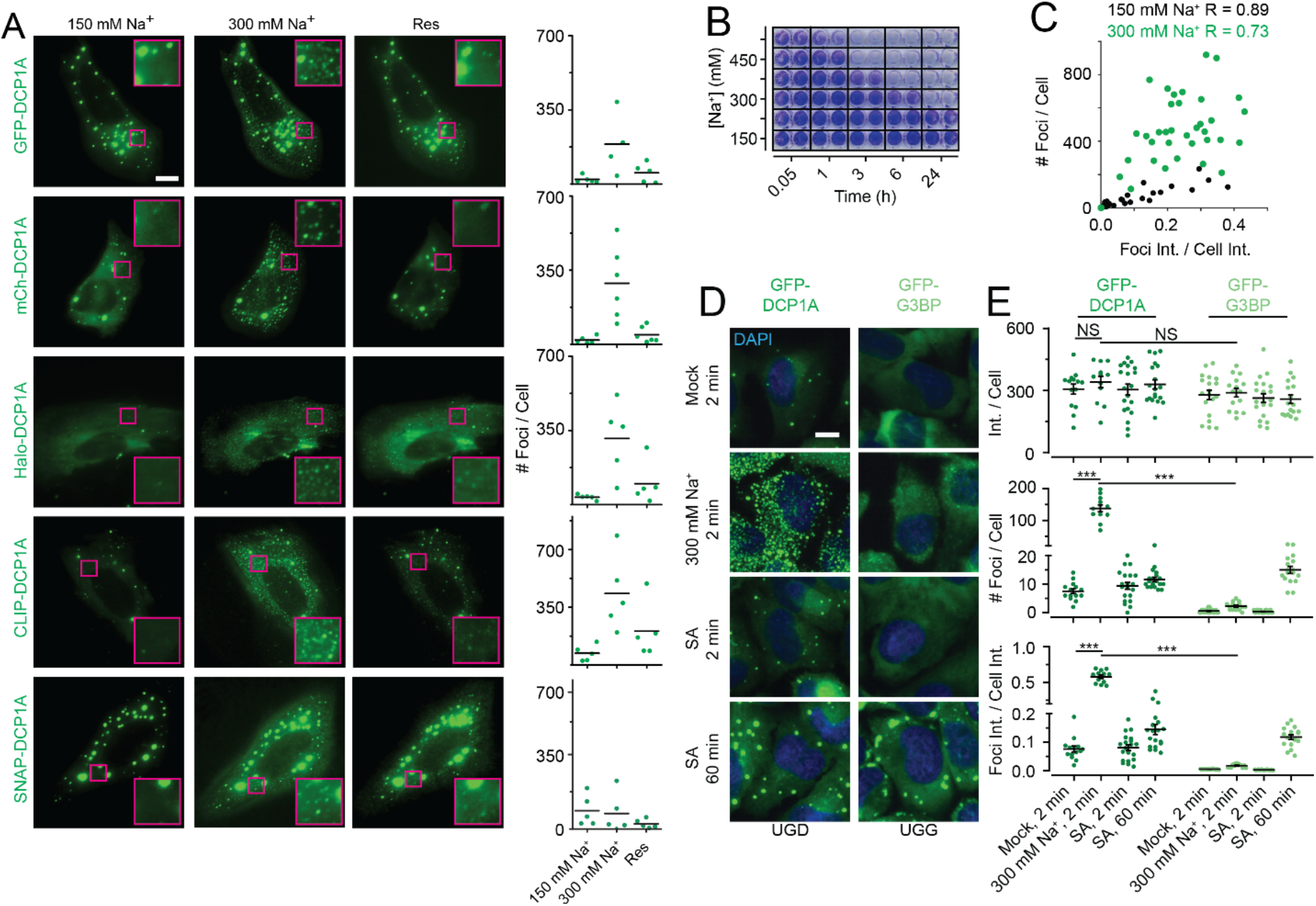
Rapid and reversible condensation of DCP1A does not depend on the fusion tag, does not affect cell viability and is distinct from G3BP condensation. **Related to Figure 2.** (A) Representative pseudocolored images of U2-OS cells expressing DCP1A fused to different types of fluorescent or fluorogenic tags (green). Cells were treated with isotonic (150 mM Na^+^, 2 min) medium, hypertonic (300 mM Na^+^, 2 min) medium, or rescued with isotonic medium (2 min) after hypertonic treatment (2 min). Scale bar, 10 µm. Scatter plot of the number of foci per cell for each treatment condition is also shown. n = 2, > 5 cells per sample. (B) Representative image of a 96-well plate probed for cell viability by crystal violet staining across various Na^+^ concentrations and multiple time points. n = 3, with technical replicates for each n. (C) Correlation plot of the number of foci per cell against the partition coefficient in the same cell. R = correlation coefficient. Black data points = 150 mM Na^+^ treatment, green data points = 300 mM Na^+^ treatment. (D) Representative pseudocolored images of cells stably expressing GFP-DCP1A (UGD, green) or GFP-G3BP (UGG, green) and subjected to osmotic or SA stress for various amounts of time. After fixation, cells were stained with DAPI (blue). Scale bar, 10 µm. (E) Scatter plots of area normalized cell intensity (top), number of foci per cell (middle) and partition coefficient (bottom). Each dot represents a cell. Line, mean, error bars, standard error. n = 3, > 15 cells per condition, NS = not significant, ***p < 0.0001, significance by two-tailed, unpaired Student’s t-test.

**Figure S3.**
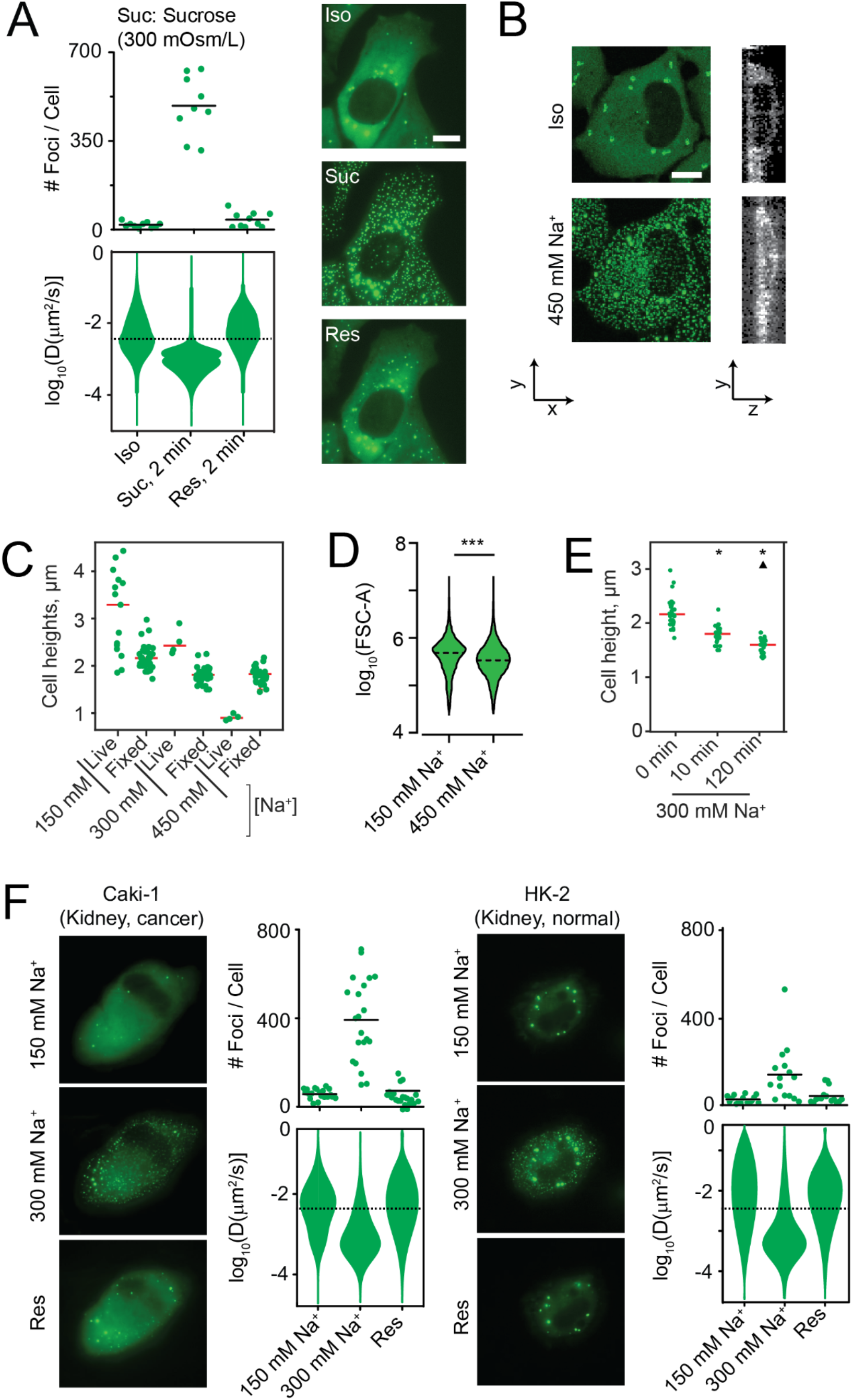
Hyperosmotic phase separation of DCP1A is correlated with cellular compression and is independent of cell type. **Related to Figure 3.** (A) Scatter plot of the number of foci per cell (top), violin plots of diffusion constants associated with DCP1A foci (bottom). Representative pseudocolored images of UGD cells (GFP, green) were treated with isosmotic (Iso) growth medium, hyperosmotic growth medium containing the non-ionic osmolyte Sucrose (Suc, 2min) or rescued (Res) with isosmotic medium (2 min) after sucrose treatment (2 min). n = 2, > 5 cells per sample. Scale bar, 10 µm. (B) Representative x-y (green) and y-z (gray) projection of a UGD cell from 3-D imaging assay wherein the cell was treated with isotonic (150 mM Na^+^) medium or hypertonic (300 mM Na^+^) medium. n = 1, 4 cells per sample. Scale bars, 10 µm (x and y) and 5 µm (z). (C) Scatter plots representing cell heights measured by DiI staining and 3-D imaging in live- and fixed-cells. Cell heights were measured from YZ sections of Z-stack images of UGD. n=2, >8 cells per replicate. (D) Violin plots of forward scatter from fixed UGD cells as measured by flow-cytometry. Cells were treated with isotonic (150 mM Na^+^) or hypertonic (300 mM Na^+^) medium for 2 min. n = 3, > 50,000 events per condition, ***p < 0.0001, significance by two-tailed, unpaired Student’s t-test. Dotted line, median. (E) Scatter plots representing cell heights measured by DiI staining and 3-D imaging in fixed-cells. Cells were treated with hypertonic (300 mM Na^+^) medium prior to fixation. Cell heights were measured from YZ sections of Z-stack images of UGD, n=2, >8 cells per trial. (F) Representative pseudocolored images of Caki-1 or HK-2 cells expressing GFP-DCP1A (green). Cells were treated with isotonic (150 mM Na^+^, 2 min) medium, hypertonic (300 mM Na^+^, 2 min) medium or rescued with isotonic medium (2 min) after hypertonic treatment (2 min). Scale bar, 10 µm. Scatter plot of the number of foci per cell (top) and violin plots of diffusion constants associated with DCP1A foci (bottom) for each treatment condition for Caki-1 or HK-2 cells are also shown. n = 2, > 5 cells per sample.

**Figure S4.**
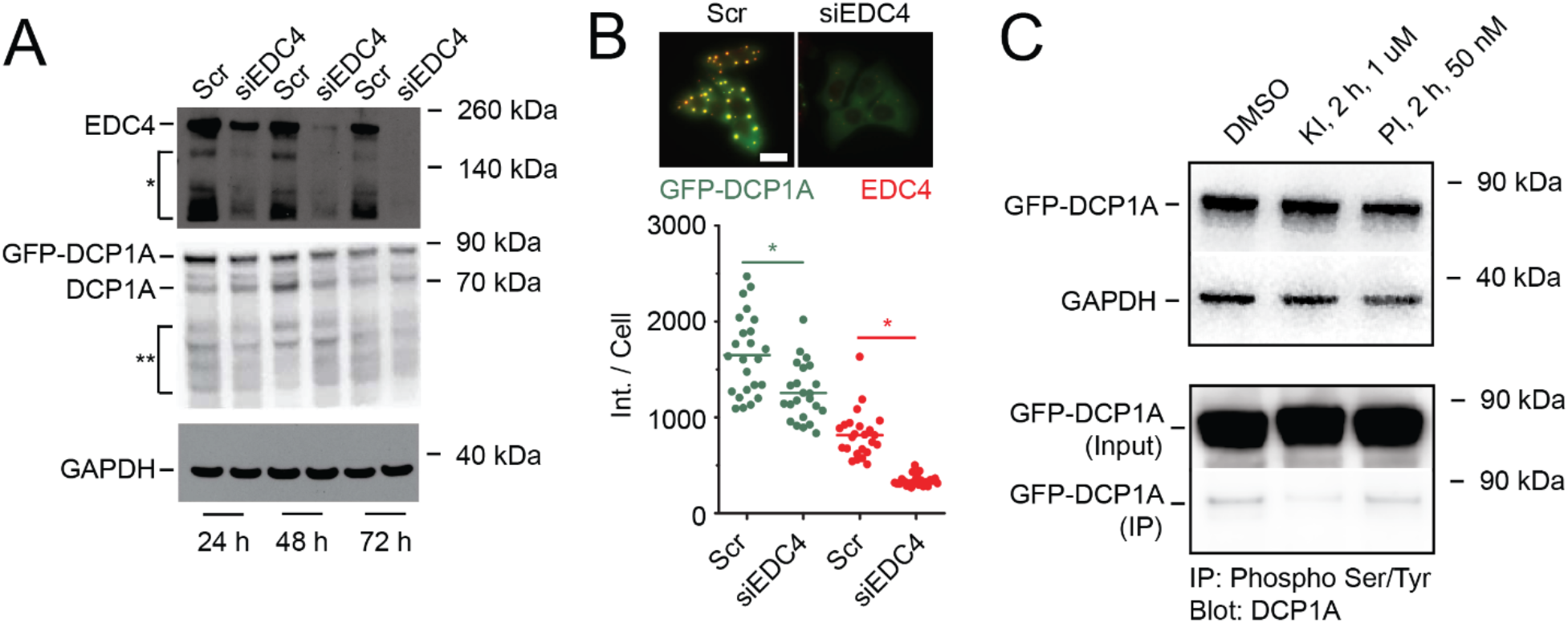
Knockdown of EDC4 results in reduced expression of DCP1A, but kinase inhibitor treatment only reduced DCP1A phosphorylation. **Related to Figure 4.** (A) Western Blot of EDC4, DCP1A, and GAPDH after various siRNA treatment times (24, 48 and 72 hr post siRNA transfection). Bands labeled with “*” and “**” were detected by EDC4 and DCP1A antibodies respectively and either denote non-specific bands or shorter protein fragments. (B) Representative pseudocolored IF images of UGD cells (top) expressing GFP-DCP1A (green), stained for EDC4 (red). Scale bar, 20 µm. Cells were either transfected with a scrambled siRNA (Scr) or siEDC4 for 48 h. Scatter plot (bottom) of the average intensity of GFP (green) or EDC4 (Cy5, red) per UGD cell transfected with a scrambled siRNA (Scr) or siEDC4 in isotonic conditions. n = 2, > 20 cells per sample, *p ≤ 0.01, by two-tailed, unpaired Student’s t-test. (C) Western blot of DCP1A, and GAPDH after various drug perturbations (top). Western blot of DCP1A after immunoprecipitation with phospho-specific antibody, upon various drug perturbations (bottom).

**Figure S5.**
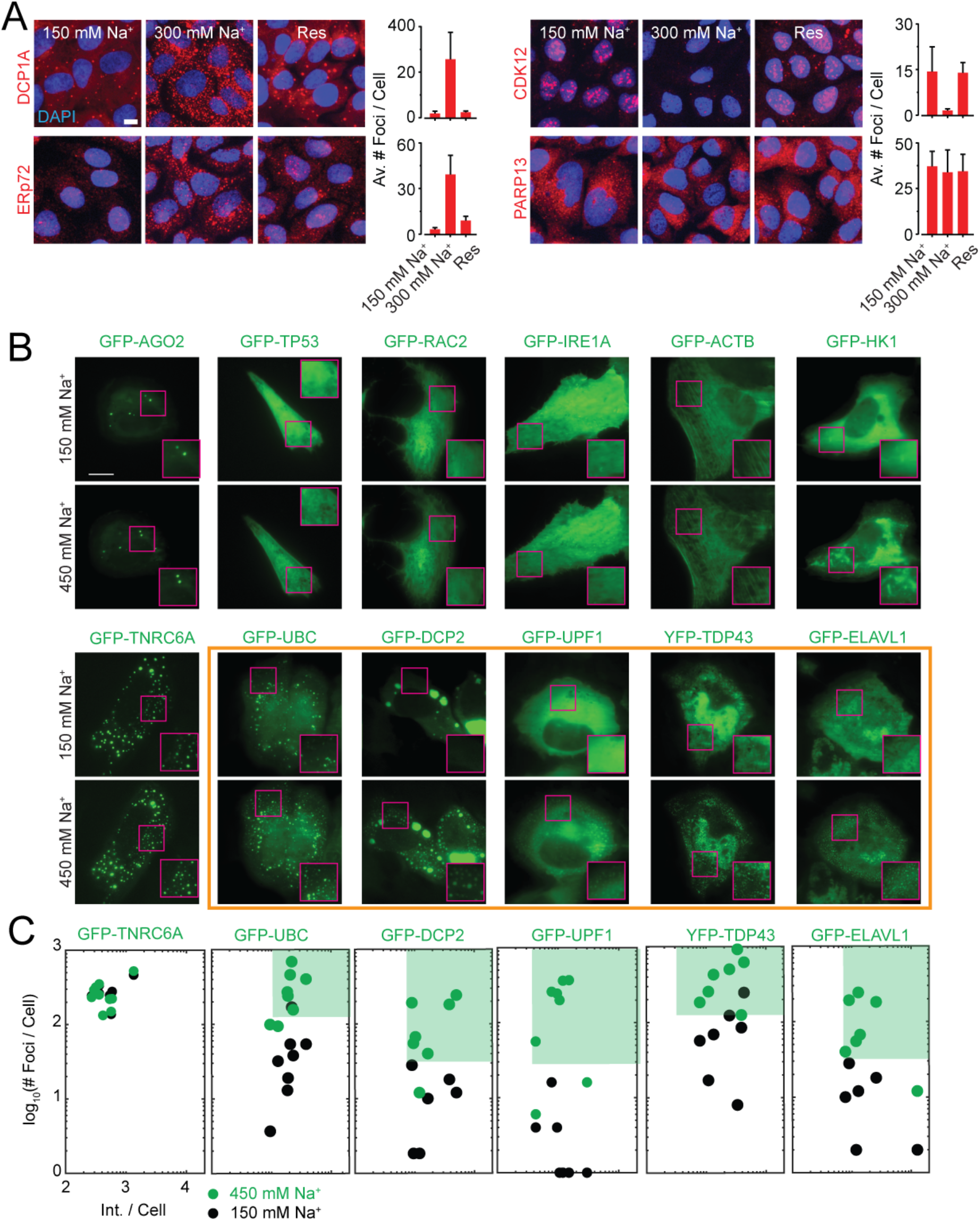
High-thoughput IF and GFP imaging of proteins in U2OS cells. **Related to Figure 5.** (A) Representative pseudocolored images of U2OS cells stained with the appropriate antibody (red, DCP1A, ERp72, CDK12, PARP13). Cells were treated with isotonic (150 mM Na^+^) medium (2 min), hypertonic (300 mM Na^+^) medium (2 min) or rescued (res) with isotonic medium (2 min) after hypertonic treatment (2 min). Scale bar, 10 µm. Quantification of the average number of foci per cell is also depicted. Error bar, standard deviation. n = 3, > 50 cells. (B) Representative pseudocolored images of U2OS cells (GFP, green) transfected with the appropriate GFP-tagged construct and treated with isotonic (150 mM Na^+^) medium or hypertonic (300 mM Na^+^) medium for 2 min. Scale bar, 10 µm. Inset depicts a zoomed-in area corresponding to a 15 x 15 µm^2^ magenta box. Constructs that exhibit HOPS are highlighted in orange. (C) Scatter plot of the number of foci per cell against the area-normalized total cell fluorescence intensity for each GFP-labeled protein. n = 2, > 5 cells per sample. The green contour depicts HOPS.

**Figure S6.**
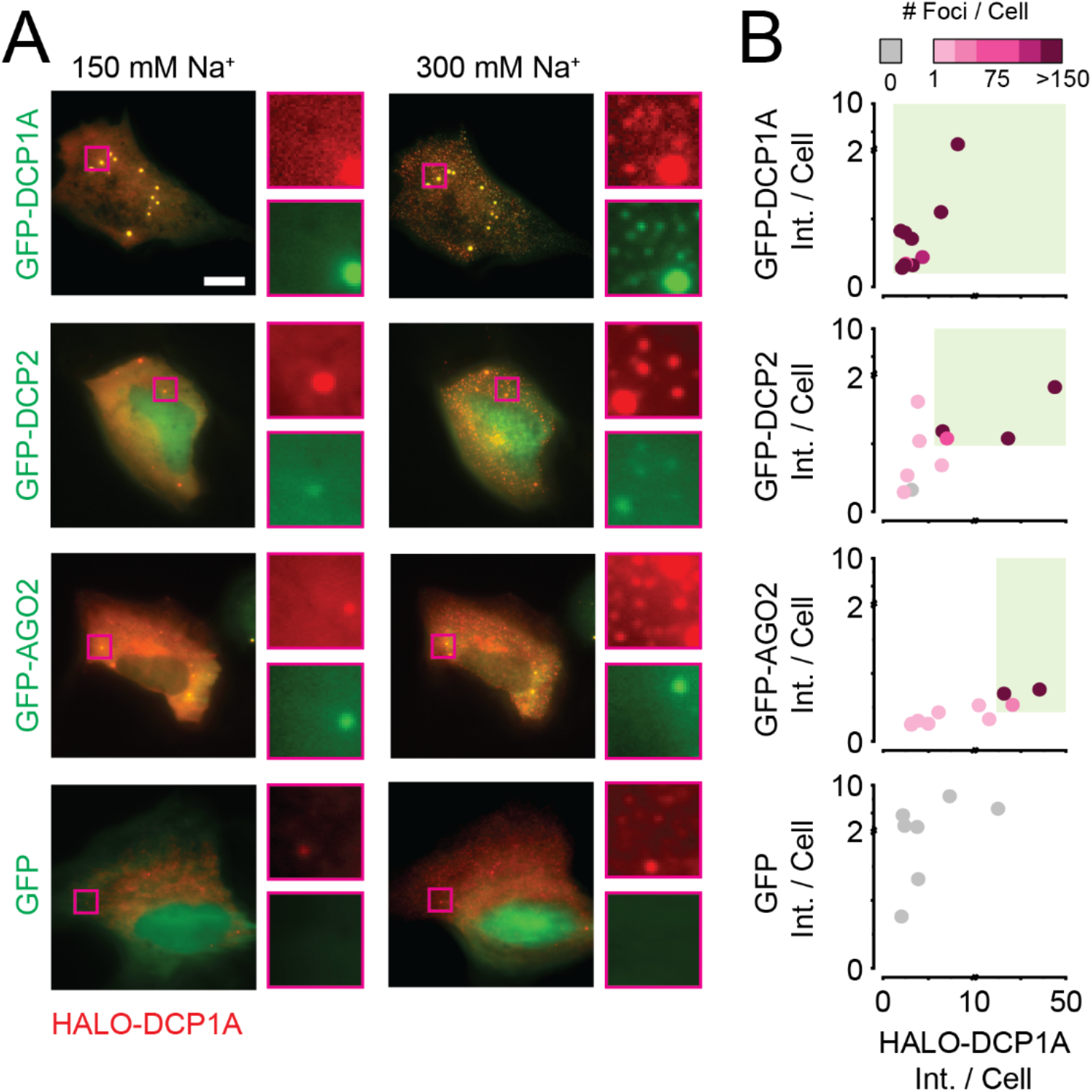
Interactors of DCP1A can exhibit HOPS at high DCP1A concentration. **Related to Figure 5.** (A,) Representative pseudocolored images of U2OS cells (GFP, green, JF646, red) transfected with GFP-tagged DCP1A (positive control), DCP2, AGO2, or GFP (negative control) and treated with isotonic (150 mM Na^+^) or hypertonic (300 mM Na^+^) medium for 2 min. Scale bar, 10 µm. Insets depict magnified areas corresponding to a 2 x 2 µm^2^ box as indicated; n = 2, 5 cells per replicate. (B) Color mapped scatter plots of area normalized Halo-DCP1A fluorescence intensity against area normalized GFP-tagged protein intensity. Each dot represents a cell and its color represents the number of foci within the cell upon hypertonic (300 mM Na^+^) treatment. The color scheme is provided on the top of the plot. Green contours depict HOPS conditions, which is defined by an at least 3-fold increase in the number of foci under hypertonic conditions compared to isotonic controls.

**Figure S7.**
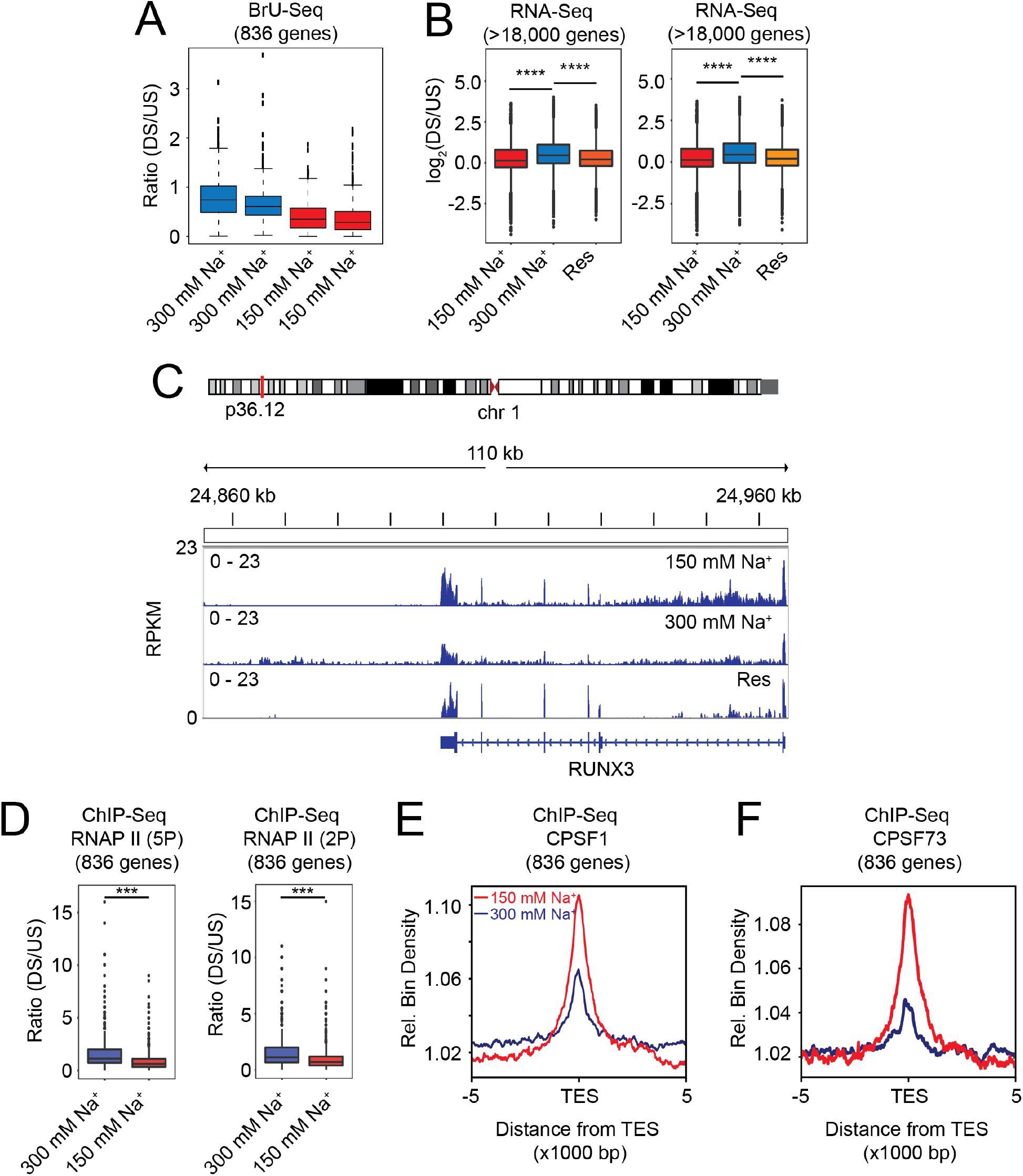
Hyperosmolarity-induced transcript read-through correlates with loss of TES occupancy by CPSFs. **Related to Figure 6.** (A) Ratio between read counts downstream (DS) and read-counts upstream (US) of TES for 836 genes assayed by BrU-Seq for each replicate. Cells were treated with isotonic (150 mM Na^+^, 30 min) or hypertonic (300 mM Na^+^, 30 min) mediums prior to sequencing. (B) DS:US ratio of > 18,000 genes that show transcript read-through in RNA-Seq assays. Cells were treated with isotonic (150 mM Na^+^, 4 h) medium, or hypertonic (300 mM Na^+^, 4 h) medium, or rescued (Res) with isotonic medium (4 h) after hypertonic treatment (4 h) prior to sequencing. (C) RNA-seq tracks of the RUNX3 locus under isotonic (150 mM Na^+^, 4 h) medium, hypertonic (300 mM Na^+^, 4 h) medium, or rescued (Res) with isotonic medium (4 h) after hypertonic treatment (4 h) prior to sequencing. (D) Ratio between ChIP-seq read counts downstream (DS) and upstream (US) of TES for 836 genes. Left panel, ChIP-seq with antibody against Serine 5P in C-terminal domain (CTD) of RNA Pol II; right panel, ChIP-seq with antibody against Serine 2P in CTD of RNA Pol II. Cells were treated with isotonic (150 mM Na^+^, 30 min) or hypertonic (300 mM Na^+^, 30 min) medium prior to ChIP-seq. (E and F) Aggregated ChIP-seq reads of CPSF1 (E) and CPSF73 (F) for 836 genes mapped around the TES. Cells were treated with isotonic (150 mM Na^+^, 30 min, red) or hypertonic (300 mM Na^+^, 30 min, blue) medium prior to ChIP-seq.

**Supplementary Movie 1. Hyperosmolarity-induced DCP1A condensation is rapid and reversible.**

Video of two UGD cells undergoing hypertonic compression and recovery in real time. Cells are initially in isotonic medium which is aspirated and replaced with hypertonic medium (30-90 s, orange) which is then replaced with isotonic medium (110 s, light grey). Inset plot represents per-frame foci number over time. Arrows represent medium addition or aspiration. Dark grey portion represents changes in focal plane from physical perturbation during pipetting.

## SUPPLEMENTARY TABLES

Table S1. Targets probed by high-throughput IF. Information about each target’s annotated multimerization potential (from the Gene Ontology and individual references), localization (from the human protein atlas and our study), HOPS propensity, and the antibodies utilized are included. Related to Figure 5.

Table S2. Targets probed by GFP imaging. Information about each target’s annotated multimerization potential (from the Gene Ontology and individual references), HOPS propensity, and the expression plasmid utilized are also included. Related to Figure 5.

### KEY RESOURCE TABLE

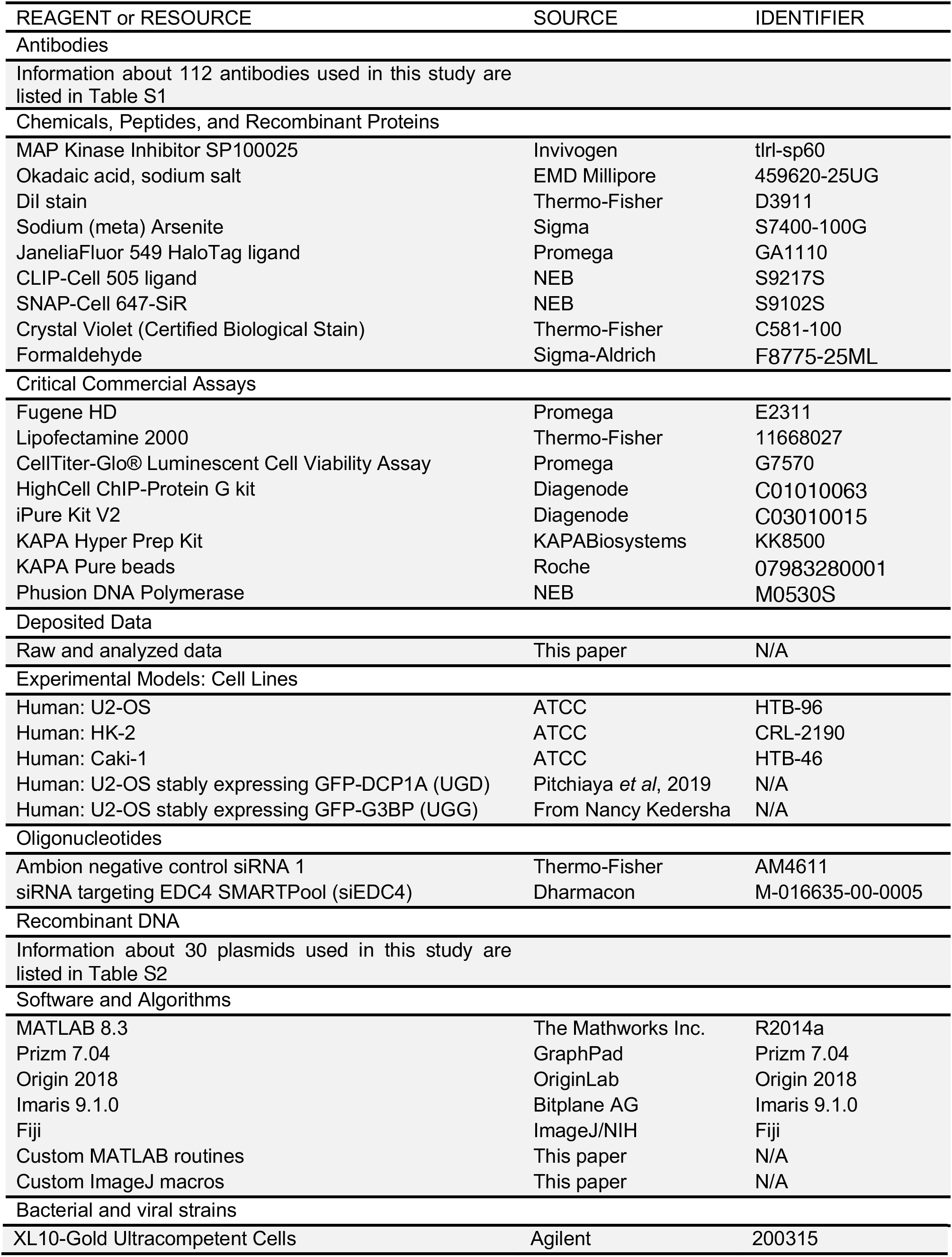

## METHODS

### DNA and RNA oligonucleotides

DNA oligonucleotide with 30 consecutive T’s (Oligo-dT36-Cy5) contained a Cy5 dye at the 3’ end and was purchased from IDT. Dyes were attached after oligonucleotide synthesis to a 3’ amino group on a C6 carbon linker and were HPLC purified by the vendor. Negative control siRNA (Scr, Ambion negative control siRNA 1) and siRNA against EDC4 (siEDC4, siRNA targeting EDC4 SMARTPool) were purchased as ready-to-use duplexes from Thermo-Fisher and Dharmacon respectively.

### Plasmids

Most plasmids were purchased from Addgene or were shared by independent labs. GFP-tagged proteins candidates were selected based on gene ontology annotation containing terms “identical protein binding” (GO:0042802), “protein homotrimerization” (GO:0070207), “protein trimerization” (GO:0070206), “protein dimerization” (GO:0046983) and “protein tetramerization” (GO:0051262) and independently verified for their self-interacting ability by the tool SLIPPER (http://lidong.ncpsb.org/slipper/index_1.html). The resulting pools of proteins were selected to cover a range of valences. The proteins tested in each class are: monomeric, p53; dimeric, AKT, Rac2; trimeric, HSF1; tetrameric, PKM2; octomeric, PAICS; IDR-containing: FUS, TDP-43. pcDH-Halo-DCP1A, pcDH-SNAP-DCP1A, pcDH-GFP-DCP1A, pcDH-mCherry-DCP1A, and pcDH-CLIP-DCP1A were constructed by first sub-cloning the DCP1A open-reading frame (ORF) from pEGFP-DCP1A into the pcDH backbone to generate pcDH-DCP1A. The ORFs of Halo, SNAP, GFP, mCherry and CLIP were PCR amplified from pFN21A (Promega), pSNAPf (NEB), pEGFP-C1 (Clontech), pEF1a-mCherry (Clontech), and pCLIPf (NEB), respectively. These amplicons were then sub-cloned into the pcDH-DCP1A backbone to generate the appropriate plasmids.

### Cell culture

U2OS and U2OS-GFP-DCP1A (UGD) cells were propagated in McCoy’s 5A medium supplemented with 10% fetal bovine serum and Penicillin-Streptomycin (GIBCO, #15140). UGD cells were kept under positive selection with 100 µg/mL G418. Hypertonic medium was prepared by supplementing regular growth medium with 10x PBS such that the appropriate sodium concentration was achieved. Isotonic medium was prepared by mock supplementing regular growth medium with 1x PBS, whose volumes matched that of 10x PBS in hypertonic medium. Oxidative stress was induced by treating cells with 0.5 mM sodium arsenite (SA). Hyperosmotic medium with sucrose or sorbitol were prepared by directly dissolving the appropriate reagent to achieve 300mOsm/L (300 mM). Plasmid transfections for GFP imaging were achieved using Fugene HD (Promega, # E2311) as per the manufacturer’s protocol. UGD cells were transfected with siRNAs using Lipofectamine RNAimax (Thermo-Fisher, # 13778030) as per the manufacturer’s protocol. For live cell imaging of Halo-DCP1A, CLIP-DCP1A, and SNAP-DCP1A, cells were treated with 100 nM of the appropriate ligand for 30 min in growth medium without phenol-red. After the treatment, cells were washed three times in phenol-red free medium and placed back in the incubator for 30 min, prior to imaging. For live cell imaging, cells were imaged in phenol-red free medium containing 1% FBS and the appropriate tonicity.

For DCP1A expression time course data (Figure S2B), U2OS cells were transfected with pGFP-DCP1A using Fugene HD. Transfected cells were imaged at 12, 24, 36, 48 and 72 hours after transfection to allow the expression level of the protein to build up. Cells were images under isotonic and hypertonic conditions at each time point to cover about 2-orders of magnitude of total GFP fluorescence intensity.

### Cell viability assays

100 µL of 10,000-20,000 cells were seeded per well of a 96 well white bottom plate or 96 well transparent plate in regular growth medium. 24 h after seeding, cells were treated with appropriate isotonic or hypertonic medium. Cell growth and viability were measured on the 96 well white bottom plate as an end point measurement for each time point and/or treatment using the Cell-titer GLO assay (Promega, # G7570) based on manufacturer’s instructions. 96 well transparent plates were fixed with 100% methanol at RT for 10 min, stained with crystal violet (0.5% in 20% methanol) for 20 min at RT, washed with water and photographed.

### Immunofluorescence

Cells were grown on 8 well chambered coverglasses (Thermo-Fisher, # 155383PK), washed with PBS, formaldehyde fixed and permeablized using 0.5% Triton-X100 (Sigma, T8787-100ML) in 1x PBS at room temperature (RT) for 10 min. It is important that the tonicity of the wash buffer and fixative matched that of the cell medium. Cells were then treated with blocking buffer containing 5% normal goat serum (Jackson Immunoresearch, 005-000-121), 0.1% Tween-20 (Sigma, P9416-50ML) in 1x PBS at RT for 1 h. Primary antibodies were diluted in blocking buffer to appropriate concentrations. Cells were incubated with primary antibodies at RT for 1 h. Following three washes with the blocking buffer for 5 min each, cells were treated with secondary antibodies diluted in blocking buffer to appropriate concentrations. Then, following two washes with the blocking buffer and two washes with 1x PBS for 5 min each, cells were mounted in solution containing 10 mM Tris/HCl pH 7.5, 2 × SSC, 2 mM trolox, 50 μM protocatechuic acid, and 50 nM protocatechuate dehydrogenase. Mounts were overlaid with mineral oil and samples were imaged immediately.

### Combined IF and RNA fluorescence *in situ* hybridization

Following the final 1x PBS washes in the abovementioned IF protocol, cells were formaldehyde fixed and permeablized overnight at 4°C using 70% ethanol. Cells were rehydrated in a solution containing 10% formamide and 2 × SSC for 5 min and then treated with 100 nM Oligo-ribonucleoside complex, 0.02% RNase-free BSA, 1 μg/μl *E. coli* transfer RNA and 10% formamide at 37 °C. After hybridization, cells were washed twice for 30 min at 37 °C using a wash buffer (10% formamide in 2 × SSC). Cells were then mounted in solution containing 10 mM Tris/HCl pH 7.5, 2 × SSC, 2 mM trolox, 50 μM protocatechiuc acid, and 50 nM protocatechuate dehydrogenase. Mounts were overlaid with mineral oil and samples were imaged immediately.

### Microscopy

Highly inclined laminated optical sheet (HILO) imaging was performed as described (Pitchiaya et al., 2012; Pitchiaya et al., 2013, Pitchiaya et al., 2017, Pitchiaya et al., 2019) using a cell-TIRF system based on an Olympus IX81 microscope equipped with a 60x 1.49 NA oil-immersion objective (Olympus), as well as 405 nm (Coherent, 100 mW at source, ∼65 µW for imaging CB-Dex), 488 nm (Coherent ©, 100 mW at source, ∼1.2 mW for imaging GFP), 561 nm (Coherent ©, 100 mW at source, ∼50 µW for imaging mCh) and 640 nm (Coherent ©, 100 mW at source, 13.5 mW for imaging Cy5) solid-state lasers. Quad-band filter cubes consisting of z405/488/532/640rpc or z405/488/561/640rpc dichroic filters (Chroma) and z405/488/532/640m or z405/488/561/640m emission filters (Chroma) were used to filter fluorescence of the appropriate fluorophores from incident light. Emission from individual fluorophores was detected sequentially on an EMCCD camera (Andor IXon Ultra) for fixed cell imaging. For live cell imaging cells were seeded on Delta T dishes (Bioptechs, 04200417C) and imaged on a Bioptechs temperature control module (Bioptechs, 0420-4). High-throughput IF was performed on the same microscope using a 60x 0.9 NA air objective. The multi-well scanning mode in Metamorph®, the acquisition software, was used to control a motorized stage (MS-2000, Applied Scientific Instrumentation Inc.).

### Image Analysis

For measuring the average GFP signal per cell, GFP intensity thresholds were set (Huang threshold in ImageJ) to automatically identify cell boundaries. Background intensity, outside of cell boundaries, was subtracted from GFP signal to extract the background corrected GFP intensity within cells. The corrected intensity was then divided by the total number of thresholded (Huang threshold in ImageJ) DAPI stained nuclei to extract the average GFP intensity per cell. For measuring the percentage of GFP signal within foci, images were first thresholded (percentage threshold in ImageJ) to create masks of foci and the GFP intensity within this mask was calculated. Background corrected foci intensity was then divided by the background corrected GFP intensity within cells. Average number of foci per cell in IF images were identified using the find maxima function in ImageJ. Briefly, a 5-pixel radius rolling ball was used to subtract the background from images, which were subsequently convolved with a 5×5 pixel kernel and a 2-pixel radius Gaussian blur function. These image processing steps enhanced the definition of a spot that were easy to identify with the find maxima function. The noise tolerance (or threshold value) in the find maxima function was maintained across samples that were compared. The total number of spots was then divided by the number of nuclei to calculate the mean spots per cell. Imaris was used for single particle tracking. Custom Matlab scripts were used to extract diffusion rates of the trajectories by fitting the mean squared displacement (< *r*^2^ >) over the first five observation time windows to a line and extracting the slope. Diffusion rates ($) were then calculated as per the 2-D diffusion equation from

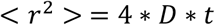

The obtained logarithm of the obtained diffusion values was plotted as histograms in Origin which were then visualized as violin plots using custom scripts in Matlab.

Intracellular GFP-DCP1A concentrations were determined using an eGFP calibration curve, obtained by capturing multi-frame movies of a range (1 nM – 30 µM) of EGFP dilutions in 1x PBS at the same camera acquisition settings used for imaging GFP-DCP1A. The average pixel intensity over 3 fields of view over 5 frames was fitted with a linear equation that was used to then assign mean intracellular GFP-DCP1A intensities to concentrations. In plots of foci number against concentration or intensity, the HOPS regime was drawn by defining the lowest concentration or intensity (x-axis limit) that showed either >100 foci or a threefold change in foci number under hypertonic treatment relative to isotonic, whichever was higher. The partition coefficient was calculated as:

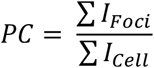

where *I_Foci_* represent total intensity in foci in a cell, and *I_Cell_* represents total intensity in the cell determined by semi-manual thresholding using custom scripts in Matlab. The phase plot in Figure 2E was fit using a dose response equation in Origin:

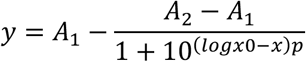

The best fit to the binned data was used to determine the PC_50_, given by log(x0). The hypertonic data were fit with p = 2 and isotonic data were fit with p = 1. All final figures were assembled in Adobe Illustrator.

### Flow cytometry

Cell volume was evaluated using a Sony SH800 cell sorter on 100 µm sorting chips sheathed with 1x PBS (Life Technologies Applied Biosystems, Vic, Australia). Trypsinized U2-OS or UGD cell suspensions were treated with isotonic (150 mM Na^+^) or hypertonic (300 mM Na^+^) growth medium for 2 min, immediately fixed in pre-warmed (37 °C) 4% paraformaldehyde solution with matched tonicity for 10 min at 25 °C and washes 3 times with 1x PBS. Upon cell loading, GFP positive UGD cells were identified by fluorescence compensation from U2-OS cells via 488 nm illumination. GFP-gated cells were then analyzed for their forward scattering extent, since it is considered to related to the volume of the cell (Model, 2018).

### RNAseq and Bru-seq

For steady-state RNAseq, UGD cells were grown in 10 cm dishes, treated with the appropriate medium (isotonic, 150 mM Na^+^ or hypertonic, 300 mM Na^+^) and cells were harvested by scraping in RIPA buffer (Thermo-Fisher, PI89900). Total RNA was then extracted with QIAGEN RNeasy midi kit (Cat. No. 75144). RNA integrity was assessed using an Agilent Bioanalyzer. Each sample was sequenced in duplicated using the Illumina HiSeq 2000 (with a 100-nt read length). Strand-specific paired-end reads were then inspected for sequencing and data quality (for example, insert size, sequencing adaptor contamination, rRNA content, sequencing error rate). Libraries passing quality control were trimmed of sequencing adaptors and aligned to the human reference genome, GRCh38. Sample were demultiplexed into paired-end reads using Illumina’s bcl2fastq conversion software v2.20. Reads were mapped onto hg38 human reference genome using TopHat2. First the reference genome was indexed using bowtie2-build. Paired end reads were then aligned to the reference genome using TopHat2 with strand-specificity and allowing only for the best match for each read. Aligned file was used to calculate strand specific read count for each gene using bedtools multicov with -s option. A known genes gtf file downloaded from UCSC was used to calculate read count. Two additional bed files were created for each gene representing 10 kb upstream and 10 kb downstream of the TSS. For each gene, read count was calculated for its upstream and downstream region as well with strand-specificity. To estimate an RNA read-through event, we calculated the ratio of read count for 10kb downstream of TSS to 10kb upstream of TSS after normalizing it for gene expression and sequencing depth. A box plot was plotted for this normalized ratio for the three samples using R software and ggplot2 package. Evaluation of significance was performed using the student’s t-test. The aligned bam file of the sample was converted into bigwig format using deepTools bamcoverage. The resultant bigwig file was uploaded onto IGV for viewing of the RNA read-through event.

For Bru-seq, UGD cells were grown on T75 flasks to >80% confluency. Flasks were washed once with fresh medium before bromouridine (BrU) treatment. BrU solution was diluted to a final concentration of 2 mM in McCoy’s 5A medium containing 2% FBS containing 145 mM (isotonic) or 300 mM (hypertonic) monovalents. Cells were incubated in the appropriate bromouridine-containing medium for 30 min. Cells treated with hypertonic medium were allowed to recover in isotonic medium for 30 min or 6 h in isotonic medium before they were harvested. Nascent transcript libraries for Bru- and Bru-Chase seq were performed and sequenced as described (Paulsen et al, 2014).

Data from both RNAseq and Bru-seq were analyzed as follows. We identified the transcription end sites (TES) of genes by GENCODE annotation and defined a 10 kb region upstream (US) and downstream (DS) of each TES, especially for genes that did not have any neighboring gene DS within the 10 kb distance. We then computed reads per kilobase million (rpkm) values for these US and DS bins and computed a DS/US ratio.

### Chromatin Immunoprecipitation followed by sequencing (ChIP-seq) and analysis

ChIP-seq experiments were carried out using the HighCell# ChIP-Protein G kit (Diagenode, Cat#: C01010063) following the manufacturer’s protocol. Chromatin from five million cells was used per ChIP reaction with 6.5 μg of the target protein antibody. In brief, cells were exposed to isotonic (150 mM Na^+^) or hypertonic (300 mM Na^+^) conditions for 30 min, trypsinized in solutions that maintained tonicity and washed twice with 1x PBS (isotonic) or 2 x PBS (hypertonic). This was followed by cross-linking for 12 min in 1% formaldehyde solution in iso- or hyper-osmotic PBS (Sigma-Aldrich, Cat#: F8775-25ML). Crosslinking was terminated by the addition of 1/10 volume 1.25 M glycine for 5 min at room temperature. This was followed by cell lysis and sonication (Bioruptor, Diagenode) resulting in an average chromatin fragment size of 200 bp. Fragmented chromatin was used for immunoprecipitation using various antibodies with an overnight incubation at 4 °C. Immunoprecipitated DNA was de-crosslinked and purified using the iPure Kit V2 (Diagenode, Cat#: C03010015) using the manufacturer’s protocol. Purified DNA was quantified using Qubit-3 (Invitrogen) and 10-100 ng of total DNA was used for library preparation using the KAPA Hyper Prep kit (KAPABiosystems, Cat#: KK8500) as per the manufacturer’s protocol. Briefly, DNA was first converted to blunt-ended fragments via end-repair and a single ‘A’ nucleotide was added to fragment ends. This was immediately followed by ligation of Illumina adaptors and PCR amplification (for 9-12 cycles) using Illumina barcoding primers and Phusion DNA polymerase (NEB, Cat# M0530S). PCR products were double size-selected using the KAPA Pure beads (Roche, Cat#: 07983280001) to remove fragments larger than 500 bp (0.65X beads:sample ratio) and primer dimers (0.9X beads:sample ratio). In the end, all libraries were quantified and quality checked using the Bioanalyzer 2100 (Agilent) and sequenced on the Illumina HiSeq 2500 Sequencer (paired-end, 50-nucleotide read length) to yield roughly 30-40 million total reads.

ChIP-Seq reads were aligned to the human reference genome (GRC h38/hg38) with the Burrows-Wheeler Aligner (BWA 0.7.17-r1198) (Li and Durbin, 2009). The SAM file obtained after aliment was converted into BAM format using SAMTools (Version: 1.9) (Li et al., 2009). Using deepTools bamCoverage (Ramirez et al.), a coverage file (bigWig format) for each sample was created. The coverage is calculated as the number of reads per bin, where bins are short consecutive counting windows. While creating the coverage file, the data was normalized with respect to each library size. DeepTools computeMatrix was used to calculate scores per genome regions. We used PolyASite 2.0 (Herrmann et al., 2020) to determine the locations of polyA sites on terminal exons across the human genome. These locations were used as the genome regions while running computeMatrix and calculating scores for each sample across these sites and a 1-kb window up- and downstream from these regions. To create a profile plot for scores over these polyA sites, we used deepTools plotProfile. The score matrix generated using computeMatrix was the input for plotting the score profile.

### Statistical analysis

Graphpad-Prizm and Origin were used for statistical analysis and plotting. For pairwise comparisons, p-values were calculated based on non-parametric unpaired t-tests with a Kolmogorov-Smirnov test. For comparisons involving more than 2 samples, one-way-ANOVA tests were used with a Geisser-Greenhouse correction.

